# Adipogenin Dictates Adipose Tissue Expansion by Facilitating the Assembly of a Dodecameric Seipin Complex

**DOI:** 10.1101/2024.07.25.605195

**Authors:** Chao Li, Xue-Nan Sun, Jan-Bernd Funcke, Lauri Vanharanta, Nolwenn Joffin, Yan Li, Xavier Prasanna, Megan Paredes, Chanmin Joung, Ruth Gordillo, Csaba Vörös, Waldemar Kulig, Leon Straub, Shuiwei Chen, Joselin Velasco, Ayanna Cobb, Davide La Padula, May-Yun Wang, Toshiharu Onodera, Oleg Varlamov, Yang Li, Chen Liu, Andrea R. Nawrocki, Shangang Zhao, Da Young Oh, Zhao V. Wang, Joel M. Goodman, R. Max Wynn, Ilpo Vattulainen, Yan Han, Elina Ikonen, Philipp E. Scherer

## Abstract

Adipogenin (Adig) is an evolutionarily conserved microprotein and is highly expressed in adipose tissues and testis. Here, we identify Adig as a critical regulator for lipid droplet formation in adipocytes. We determine that Adig interacts directly with seipin, leading to the formation of a rigid complex. We solve the structure of the seipin/Adig complex by Cryo-EM at 2.98Å overall resolution. Surprisingly, seipin can form two unique oligomers, undecamers and dodecamers. Adig selectively binds to the dodecameric seipin complex. We further find that Adig promotes seipin assembly by stabilizing and bridging adjacent seipin subunits. Functionally, Adig plays a key role in generating lipid droplets in adipocytes. In mice, inducible overexpression of Adig in adipocytes substantially increases fat mass, with enlarged lipid droplets. It also elevates thermogenesis during cold exposure. In contrast, inducible adipocyte-specific Adig knockout mice manifest aberrant lipid droplet formation in brown adipose tissues and impaired cold tolerance.

## Introduction

Adipogenin (Adig), comprised of 80 amino acids, was first detected in steatotic liver upon PPARγ1 overexpression in mice (*1*). Two independent groups demonstrated that it is highly expressed in adipose tissue and significantly upregulated during adipogenesis (*2-4*). Interestingly, a recent genome-wide association study (GWAS) showed that the human *ADIG* gene is associated with body mass index (BMI)-adjusted leptin levels (*5*). Phenotypic characterization of whole-body Adig knockout mice suggested Adig may play a role in adipose tissues (*6*). However, contradictory results have emerged regarding its intracellular localization and its role in adipogenesis (*2-4, 6*). Moreover, the molecular basis for Adig function remains largely unknown.

Seipin is a key protein in the formation of lipid droplets (LDs) (*7, 8*). Genetic studies in yeast (*9, 10*), worms (*11*), flies (*12*), mice (*13, 14*) and humans (*15*) have unambiguously revealed an essential and conserved role of seipin in lipid storage. Moreover, compelling evidence demonstrates the growth of LDs is directly associated with the presence of seipin foci, and the elimination of seipin results in aberrations of LD size and number (*16-18*). However, the precise mechanistic basis by which seipin generates LDs is still unknown. Recently, Arlt *et al.* proposed a fascinating model to explain the process of LD growth and budding based on the cryogenic-electron microscopy (Cryo-EM) structure of yeast seipin. In this model, the yeast decameric seipin alternates between two conformations within the transmembrane domains during the formation of LDs (*19*). However, it remains unknown how seipin functions in mammalian cells, as we lack the structure of the seipin transmembrane domains. Furthermore, the current yeast model requires the seipin complex to have an even number of oligomers, which is incompatible with the reported undecameric seipin structure in humans (*20*).

Here, we report a surprising role of Adig in the assembly of seipin oligomers. We find that mouse recombinant seipin complexes can form two distinct oligomers, 11-mers and 12-mers. Adig selectively stabilizes and binds to the transmembrane domains of the dodecameric seipin complex. Functionally, Adig deletion *in vitro* shows aberrant lipid droplet morphology during adipogenesis. Adig overexpression significantly enhances the growth of lipid droplets. In addition, gain and loss-of-function studies in mice reveal that Adig promotes the expansion of adipose tissue mass.

## Results

### Adig is highly expressed in adipocytes and localized in the ER

Our interest in Adig was initiated when its expression was reported in steatotic livers (*1*). Utilizing Northern blots, we independently found that *Adig* transcripts were highly enriched in adipose tissues and testis (**Figure S1A**) and were strongly induced during adipogenesis (**Figure S1B**). We then raised antibodies against Adig to determine protein expression. Consistent with the results from the Northern blots, Adig protein was highly expressed in adipose tissues and testis (**Figure 1A**). Adig protein was dramatically upregulated during adipogenesis in both brown (**Figure 1B**) and white adipocytes (**Figure S1C**), concomitant with the induction of PPARγ and Perilipin1. Importantly, ChIP-seq databases show PPARγ directly binds to the promoter region of *ADIG* in adipocytes (**Figure 1C**). The PPARγ agonist rosiglitazone induces the expression of Adig (**Figure S1D**). These results reveal that PPARγ transcriptionally controls the expression of Adig, thereby leading to high Adig protein expression in adipose tissues.

**Figure 1.**
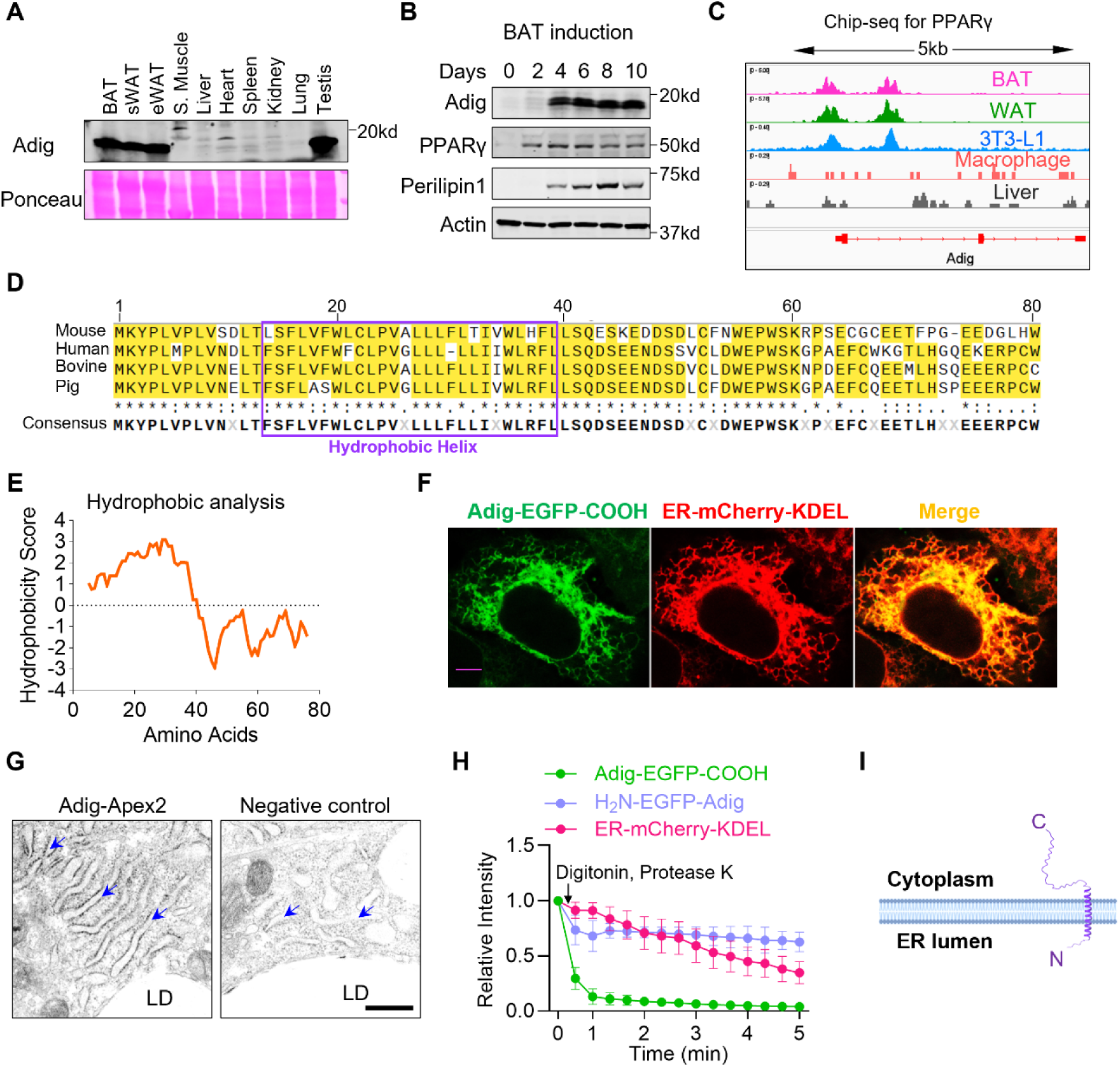
Adipogenin is highly expressed in adipocytes and located in the ER. A. The protein expression profile of Adipogenin (Adig) in different mouse tissues. Adig protein is enriched in adipose tissues and testis. The same amount of protein (2.7µg) from different tissues was loaded. B. Adig is strongly upregulated during adipogenesis in brown adipocytes *in vitro*. C. PPARγ binding peak in the Adig promoter region revealed by the Integrative Genomics Viewer. D. Amino acid sequence alignment of Adig from different species. The putative α-helix sequence is highlighted. E. Hydrophobicity analysis of amino acid sequence from mouse Adig. Scores are generated by Expasy ProtScale. F. Adig is co-localized with ER marker KDEL. Plasmids Adig-EGFP(C) and ER-mCherry-KDEL are transfected into HeLa cells. Scale bar: 5µm. G. Adig-Apex2 staining in the adipocyte. Preadipocytes are isolated from adipose tissue in Adig flox mice, and then induced into mature adipocytes. To make sure Adig-Apex2 is expressed at physiological levels, preadipocytes were infected with AAV-iCre and AAV-Adig-Apex2. For the negative control group, preadipocytes are infected with AAC-EGFP or AAV-iCre. Scale bar: 500nm. Blue arrows indicate the ER structure. H. Adig topology analysis in HeLa cells. Adig-EGFP(C) or EGFP(N)-Adig plasmids were transfected into HeLa cells. Low concentrations of digitonin permeabilize the plasma membrane without disrupting the ER structure. Proteinase K selectively cleaves the cytosolically accessible parts of proteins without affecting portions hidden in the lumen of the organelles. Fluorescence was recorded in the time lapse mode. I. Scheme for the Adig localization, topology, and putative structure.

We explored the subcellular localization of Adig, as previous reports claimed different locations, including plasma membranes (*2*), nuclei (*3*) and on the surface of lipid droplets (*4*). We first analyzed the amino acid sequence of Adig from different species. A long conserved hydrophobic α-helix stretch appears at the N-terminus of Adig (**Figure 1D and 1E**), which suggests that Adig is a transmembrane protein. We fused EGFP to either the N- or C-terminus of Adig, and co-transfected it with the endoplasmic reticulum (ER) maker ER-mCherry-KDEL or with the mitochondrial marker mito-mKeima (*21*) into HeLa or COS-7 cells. The localization of Adig-EGFP overlapped with that of ER-mCherry-KDEL (**Figure 1F**), but not with mito-mKeima (**Figure S1E**). In addition, low expression of EGFP-Adig also colocalizes with the ER (**Figure S1F**). To further determine the localization of Adig in adipocytes, we fused Adig to Apex2, an enzyme that can be used to visualize proteins for electron microscopy (EM) imaging (*22*). We transfected Adig-Apex2 into Adig knockout (KO) cells using adeno-associated virus (AAVs) at levels on par with the expression of endogenous Adig levels (**Figure S1G**). We found most of the Apex2 signal accumulated on the ER (**Figure 1G**). We determined the topology of Adig via a fluorescence-based protease protection assay in digitonin-treated cells (*23*). The EGFP from EGFP-Adig, but not from Adig-EGFP, is protected from digestion by protease K (**Figure 1H and S1H**), indicating the N-terminus of Adig is either embedded inside the bilayer of phospholipids or located within the ER lumen. Overall, these results demonstrate that Adig is an ER single-pass transmembrane protein with a cytosolic C-terminal tail (**Figure 1I**).

### Adig interacts with and stabilizes seipin

Micropeptides and microproteins typically form a complex with canonical proteins to modify their function (*24, 25*). To identify the Adig partner, we overexpressed EGFP or Adig-FLAG in adipocytes with AAVs, and then pulled down the Adig interactive proteins with a FLAG antibody. Samples were subsequently applied to mass spectrometry to identify interacting proteins (**Figure 2A and S2A**). Intriguingly, the most enriched protein with high coverage turned out to be seipin (**Figure 2B**, **Table S1**). Therefore, we speculated that Adig may interact with seipin. In addition, LDAF1, which forms a tight complex with seipin (*26, 27*) (**Figure 2B**), was also enriched in the Adig pull-downs.

**Figure 2.**
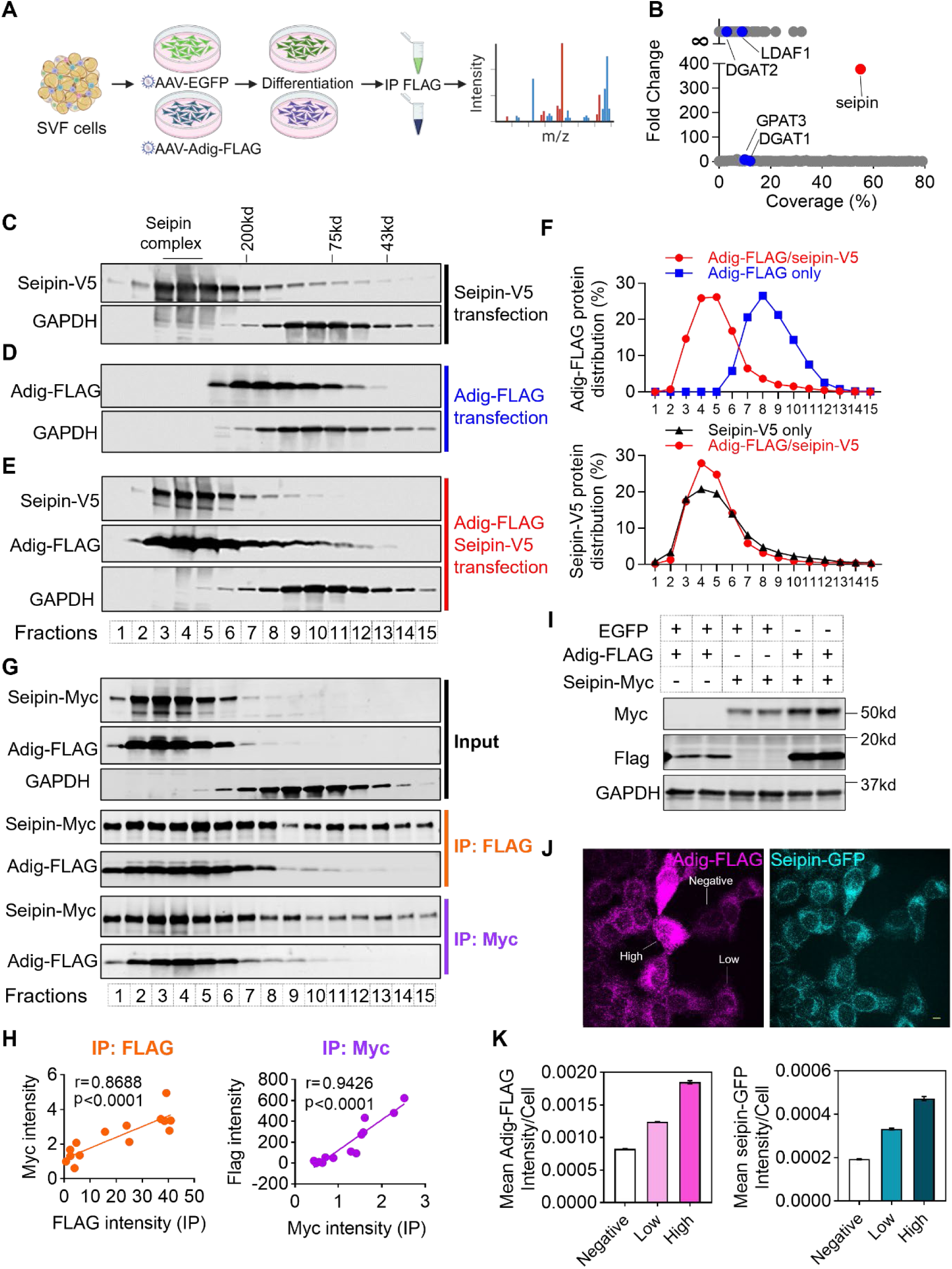
Adig and seipin form a stable complex. A. Schematic diagram of the workflow for identifying the interactive proteins of Adig. SVFs, stromal vascular fractions isolated from adipose tissues. B. Analysis for all identified proteins pulled down by FLAG immunoprecipitation. Seipin appears to be the most enriched protein in Adig-FLAG overexpression adipocytes compared to EGFP overexpression cells. C. Gel filtration analysis for cell lysis from HEK293T cells with Seipin-V5 overexpression. The HiLoad Superdex-75 column was used to fraction the elution. V5 and GAPDH antibodies were applied to Western blotting. D. Gel filtration analysis for cell lysis from HEK293T cells with Adig-FLAG overexpression. The HiLoad Superdex-75 column was used to fractionate the complex. FLAG and GAPDH antibodies were used for Western blotting. E. Gel filtration analysis for cell lysis from HEK293T cells with Seipin-V5/Adig-FLAG co-expression. The HiLoad Superdex-75 column was used to fractionate the lysate. V5, FLAG and GAPDH antibodies were applied for Western blotting. F. Distribution analysis for the Seipin-V5 and Adig-FLAG expression in fractions from (C, D and E). The expressions of V5 and FLAG were quantified by LI-COR Image Studio Lite first, and the percentage was calculated by dividing the individual expression in each fraction with the sum of all fractions combined. The peak of Adig-FLAG shifts from the lower molecular weight to the higher molecular weight fractions when Adig-FLAG is co-expressed with Seipin-V5. The Seipin-V5 accumulated more prominently in the high molecular weight fractions when Adig-FLAG was present. G. Immunoprecipitation (IP) analysis of the fractions from the gel filtration experiment. Seipin-Myc and Adig-FLAG were co-expressed in HEK293T cells, and then cells were lysed and applied to gel filtration (HiLoad Superdex-75 column). Immunoprecipitation was performed with Myc and FLAG antibodies for each fraction. H. Correlation analysis for the enriched Seipin-Myc and Adig-FLAG protein in the different fractions from (G). The expressions of Myc and FLAG were quantified by LI-COR Image Studio Lite. The Pearson correlation coefficient test was performed in GraphPad Prism. I. Western blot analysis for the expression of Seipin-Myc and Adig-FLAG in HEK293A cells. The same amount of Seipin-Myc or Adig-FLAG plasmids were transfected into the cells. J. Immunofluorescence analysis for the endogenous expression of seipin. Endogenous seipin in human epithelial A431 cells was tagged with GFP. An Adig-FLAG stable expression pool was generated. Adig-FLAG and Seipin-GFP show different expression levels. Scale bar: 10 µm. K. Quantification of the fluorescence intensity of FLAG and GFP. According to the expression of Adig-FLAG, cells were separated into three groups, Negative (n=1676), Low (n=844) and High (n=348).

Seipin exists as oligomers, and the seipin complex can be separated by size exclusion chromatography (gel filtration) (*19, 28, 29*). Therefore, to explore the physical connection between Adig and seipin within the complex, we transfected seipin-V5, Adig-FLAG and the combination of seipin-V5/Adig-FLAG into HEK293T cells, and applied the corresponding cell lysates to a gel filtration column. We analyzed the distribution of seipin and Adig in all fractions using western blotting. As predicted, the seipin complex alone accumulated in fractions with high molecular weight proteins (>200 kD) (**Figure 2C**). Interestingly, Adig, when expressed alone, fell into fractions containing medium molecular weight proteins, indicating Adig may form oligomers as well (**Figure 2D and 2F**). However, when Adig and seipin were co-expressed, they were both co-eluted into the high molecular weight fractions (**Figure 2E and 2F, Figure S2B**), indicating Adig may bind to the seipin complex. Meanwhile, we noticed the percentage of seipin in high molecular fractions somewhat increased when Adig was co-expressed (**Figure 2F**), suggesting that Adig may promote the assembly of the seipin complex. To directly examine the interactions between Adig and seipin, we performed immunoprecipitations for all the fractions obtained by gel filtration (**Figure 2G**). Interestingly, seipin in all fractions (of both high and low molecular weight) displayed strong interactions with Adig, and the relative enrichment between Adig and seipin exhibited a linear relationship when comparing these fractions (**Figure 2H**), reflecting that Adig can interact with seipin monomers as well as oligomers.

Importantly, we found that seipin/Adig co-transfection strongly increases their individual expression levels compared to individual transfections (**Figure 2I**). To assess protein stability, we inhibited protein translation with cycloheximide in single and double-transfected cells (**Figure S2C**). In the absence of seipin, the expression of Adig is sharply decreased (**Figure S2D**). Without Adig the expression of seipin tended to decrease as well (**Figure S2D**), with higher levels of both proteins present upon transfection with both expression constructs. To further explore the effect of Adig on the expression of endogenous seipin, we employed human epithelial A431 cells, in which endogenous seipin is tagged with GFP and Adig is undetectable. We then transfected Adig-FLAG into these cells and analyzed the fluorescence intensity based on GFP and FLAG staining. Cells were separated into three groups, negative, low, and high expression, according to the visible expression of Adig-FLAG (**Figure 2J**). We found that as the expression of Adig is increased, the seipin fluorescence correspondingly increases (**Figure 2K**). These results strongly indicate that seipin and Adig stabilize each other via direct interactions.

### The Cryo-EM structure of seipin/Adig complex

The structures of seipin complexes from different species have been well described (*19, 20, 28, 30*). The amino acid sequence of mouse seipin is almost identical to human seipin (**Figure S3**). Therefore, as previously done for the human seipin (*20*), we purified the mouse seipin complex from Expi293 cells with seipin and Adig co-expression (**Figure S4A-C**) and determined its structure using single-particle Cryo-EM. The two-dimensional (2D) class averages revealed two wheel-like oligomers for the seipin complex consisting of undecamers and dodecamers (**Figure 3A and 3B**). After data processing, 17,212 particles were used to reconstruct the map of the undecameric seipin complex. 32,123 particles were found as a dodecameric seipin complex (**Figure S4D**). The 11-mer map is, in fact, similar to the purified human seipin complex (*20*), but with improved resolution (3.19Å *vs.* 3.8Å) (**Figure S5A-S5C**). The inner ring-like structure, corresponding to the luminal domain of seipin, is well resolved (**Figure 3C**). However, only a small part of the transmembrane (TM) segments is visible (**Figure 3C**). The 12-mer appears to be a more rigid structure with an overall resolution of 2.98Å (**Figure S5D-S5F**). We were pleasantly surprised that the TM segments from seipin are visible in this case. In addition, extra densities were found, spanning the ER bilayer, which could not be explained by sequences from seipin (**Figure 3D**). These densities appear to be inserted between adjacent seipin monomers and were very likely contributed by the N-terminus and hydrophobic helix of Adig (**Figure 3D**). The C-terminus of Adig was invisible. In summary, we obtained two structures from the sample, the undecameric seipin complex (**Figure 3C and 3E**) and the dodecameric seipin/Adig complex (**Figure 3D and 3F**). Overall, the dodecameric seipin/Adig complex was more compact and featured a larger diameter with more hydrophobic residues in and around the wheel-like structure, leading us to postulate that the dodecamer has a higher capacity to generate lipid droplets compared to the undecameric seipin-only complex.

**Figure 3.**
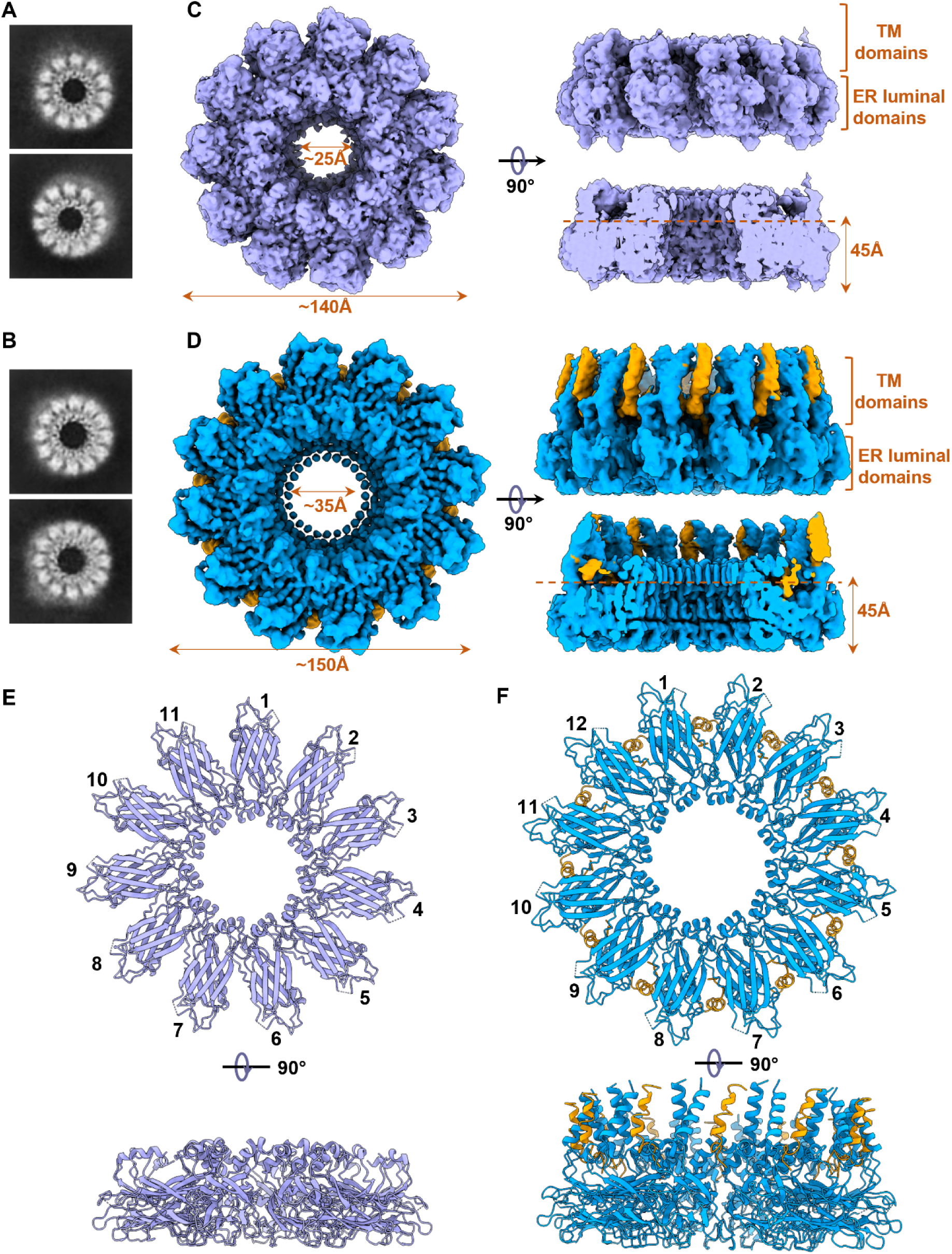
Cryo-EM structure of seipin/Adig complex. A. Representative 2D class averages for the undecameric seipin complex. B. Representative 2D class averages for the dodecameric seipin complex. C. Cryo-EM density map for the seipin complex. There is no obvious density for Adig. Top view reveals the 11 copies of the seipin subunits. The side view reveals the position of the transmembrane and luminal domains in the seipin complex. A sliced view shows the cage-like structure. D. Cryo-EM density map for the seipin/Adig complex. The Adig has a separate density from seipin and is colored brown. The top view reveals the 12 copies of seipin/Adig subunits. The side view reveals the position of the transmembrane and luminal domains in the seipin/Adig complex. The sliced view shows the cage-like structure. E. Structural model of the undecameric seipin complex. The top view is from the luminal side. F. Structural model of the dodecameric seipin/Adig complex. The top view is from the luminal side.

### Adig promotes the assembly of seipin complexes

The observation of mutual stabilization (**Figure 2**) and the structure of the seipin/Adig dodecameric complex (**Figure 3**) led us to hypothesize that Adig may promote or stabilize the assembly of seipin complexes. The enhanced assembly of seipin complexes could be very apparent when we focus on newly-synthesized seipin monomers. To test whether Adig exerts an active role in seipin assembly, we observed the assembly of new complexes upon an initial depletion of mature seipin complexes. To this end, we utilized A431 cells containing seipin-degron-GFP (*18*), in which the endogenous seipin pool can be depleted rapidly and effectively. We established a protocol in which the cells can be imaged, both in the steady state and in the recovery stage (**Figure 4A**). We found that the seipin complex foci were significantly increased in the Adig-positive cells when compared to the adjacent Adig-negative cells (**Figure 4B and 4C**). Importantly, we found that the fraction of Adig-positive seipin foci was higher in the recovery stage than in the steady state (**Figure 4D and 4E**), suggesting that newly translated seipin may have a stronger affinity for Adig.

**Figure 4.**
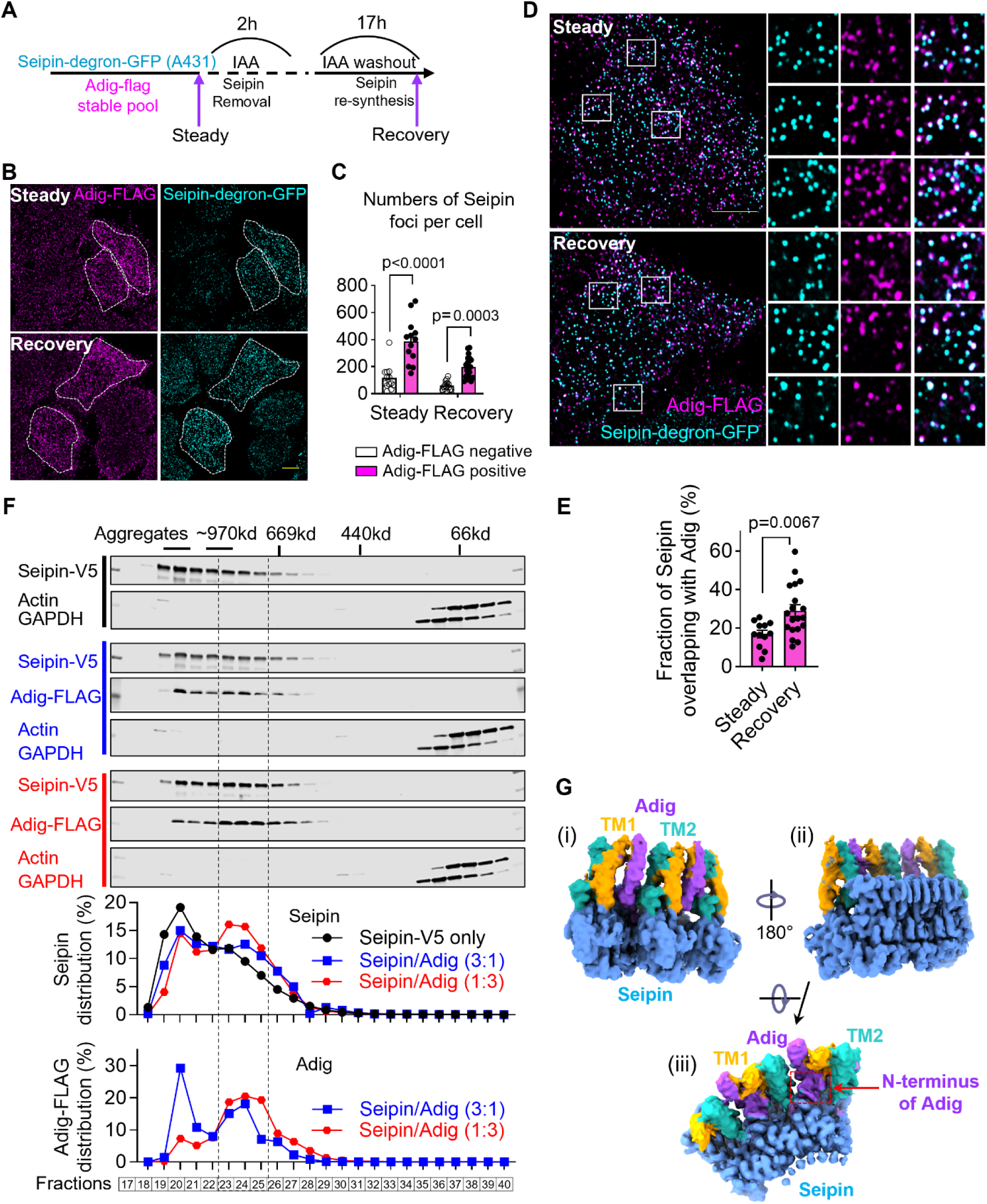
Adig promotes the assembly of seipin complexes. A. The schematic diagram shows the treatment of A431 cells. The degron and GFP sequence are knocked in the endogenous locus of the C-terminal Bscl2 gene in A431 cells. Then, the Adig-FLAG stable expression pool was generated. Seipin can be degraded by adding IAA. After the IAA was washed out, seipin can be re-synthesized. The cells were imaged at steady state (normal culture) and in the recovery state. Scale bar: 10μm. B. Adig overexpression increases the formation of seipin foci. Adig-FLAG positive cells are highlighted with a white dashed line. C. Quantification of the number of seipin foci in Adig-FLAG negative and positive cells. n=13-20. A two-way ANOVA was conducted, followed by Sidak’s multiple comparison test. Data are represented as mean±SEM. D. The colocalization between Adig-FLAG and Seipin-degron-GFP in A431 cells during the steady state and the recovery state. Scale bar: 10μm. E. Quantification for the percentage of seipin/Adig double positive foci among seipin-positive foci. In the recovery state, sepin and Adig display a higher level of co-localization. Data from 12-19 cells. Unpaired Student’s t-test was conducted. Data are represented as mean±SEM. F. Gel filtration analysis for cell lysates from HEK293T cells with Seipin-V5, Seipin-V5/Adig-FLAG (3:1) and Seipin-V5/Adig-FLAG (1:3) overexpression. A *Superose 6 Increase* column was used to fractionate the lysates. V5, FLAG, Actin and GAPDH antibodies were applied for Western blotting. The expression levels of V5 and FLAG were quantified by LI-COR Image Studio Lite first, and the percentage is calculated by dividing the individual expression level in each fraction with the total expression integrated from all fractions. Note the distribution of Seipin shifts from aggregate fractions to the native seipin complex fractions in the presence of Adig-FLAG. G. Cryo-EM map of a local region including three copies of seipin and two copies of Adig. The two transmembrane segments in seipin (TM1 and TM2) are colored brown and cyan. Adig is colored purple. (i) shows the interactions between the Adig helix and TM1/TM2. (ii) and (iii) show the position of the N-terminus of Adig.

Previous reports examined the size distribution of yeast seipin oligomers by gel filtration using the Supersoe 6 Increase resin (*19, 28*). Therefore, employing the same column (**Figure S6A**), we first tried to locate the mammalian seipin complex in different fractions. Past published data show that the overexpression of seipin and its naturally occurring mutants in mammalian cells can lead to aggregates (*31, 32*). For example, overexpression of the human mutant A212P led to ectopic large bright patches around the nuclear region, indicating a strong propensity of this mutant to form aggregates (*33*). We evaluated the distribution of overexpressed WT and naturally mutated seipin species on columns. Unexpectedly, we found that a large fraction of WT and most of the mutant seipin (A212P and L91P) migrated as aggregate-enriched fractions, compared to endogenous seipin in HeLa cells (**Figure S6B**). Endogenous seipin, the bands of which disappeared in western blots after boiling(*34*), migrated in Fractions 23-25 (**Figure S6B**). Co-expression of Adig with seipin converted seipin aggregates to the size of endogenous seipin (Fractions 23-25) (**Figure 4F**). Based on the amino acid sequences alone, the undecameric seipin complex should migrate as a ∼561 kD species, whereas the 12-mer seipin/Adig complex should migrate at ∼732 kD. With bound detergent, it is reasonable to assume that the seipin oligomers would be found in fractions ranging from 669 kD to 970 kD (**Figure 4F**). Therefore, we conclude that with the concomitant expression of Adig, seipin may tend to form normal functional seipin-Adig complexes rather than seipin aggregates.

We sought to further explore the structural basis for the effect of Adig on seipin assembly. In the map with three-seipin/two-Adig protomers, the first TM segment (TM1) of seipin and the second TM segment (TM2) from an adjacent seipin monomer may provide a hydrophobic channel that specifically accommodates the α-helix of Adig (**Figure 4G-i**). The N-terminus of Adig is embedded in the seipin oligomers, and is close to the beginning segment in the TM2 from two adjacent seipins (**Figure 4G-iii**). In addition, with the recent release of AlphaFold 3(*35*) which has provided a vast improvement in predicting the structures of the multimeric complexes, we compared the predictions of the intact architecture of seipin oligomers with or without Adig. Interestingly, we found as subunits in seipin oligomers increase from 11 to 12, the TM segments bend towards the center (**Figure S7A**). The 12-mer prediction shows that all TM segments form a cage-like structure specifically due to the interactions between Adig and seipin (**Figure S7A**). Importantly, the AlphaFold 3-generated model aligns well with our Cryo-EM map. We therefore analyzed the details of how Adig bridges two adjacent seipin monomers. Clearly, the TM segments from Adig and seipin are rich in isoleucine, leucine, and valine. Due to their hydrophobicity, these residues cluster together (**Figure S7B**). The residues in the TMs form network interactions (**Figure S7C**). The N-terminal residues in Adig interact with both the initial residues in TM2 and the curvature region in the middle of TM2 from the adjacent seipin monomer (**Figure S7D and S7E**). Therefore, we propose that there are at least two segments in Adig, namely the N-terminus and the hydrophobic helix, that mediate the interactions with seipin oligomers.

Taken together, our results strongly suggest that Adig works as a bridge to connect and stabilize the transmembrane segments of two neighboring seipin subunits. It is through these interactions that Adig promotes the assembly of the dodecameric seipin complex (**Figure S7F**). In parallel, seipin alone may spontaneously assemble into undecameric complexes as well as additional aggregates in the absence of Adig (**Figure S7F**).

### Adig facilitates the growth of lipid droplets

As seipin plays a key role at different steps of LD formation (*16-18, 26*), we explored the role of Adig in this process. In differentiated adipocytes, we determined the association of endogenous Adig and seipin by gel filtration. We found that Adig consistently co-eluted with the seipin complex (**Figure 5A**). The Adig in adipocytes migrated as a higher molecular weight complex compared to the Adig in HeLa cells with Adig-FLAG overexpression (**Figure 5A**). We generated an Adig-Apex2 construct. While we found most of the Adig-Apex2 signals were expressed on the ER (**Figure 1G**), there were also significant Apex2 positive dots in the contact sites between ER and LDs (**Figure S8A**). Furthermore, in A431 cells, Adig, seipin, and LDs displayed co-localization during LD induction (**Figure S8B**). These results suggest that seipin/Adig complexes are directly involved in LD formation.

**Figure 5.**
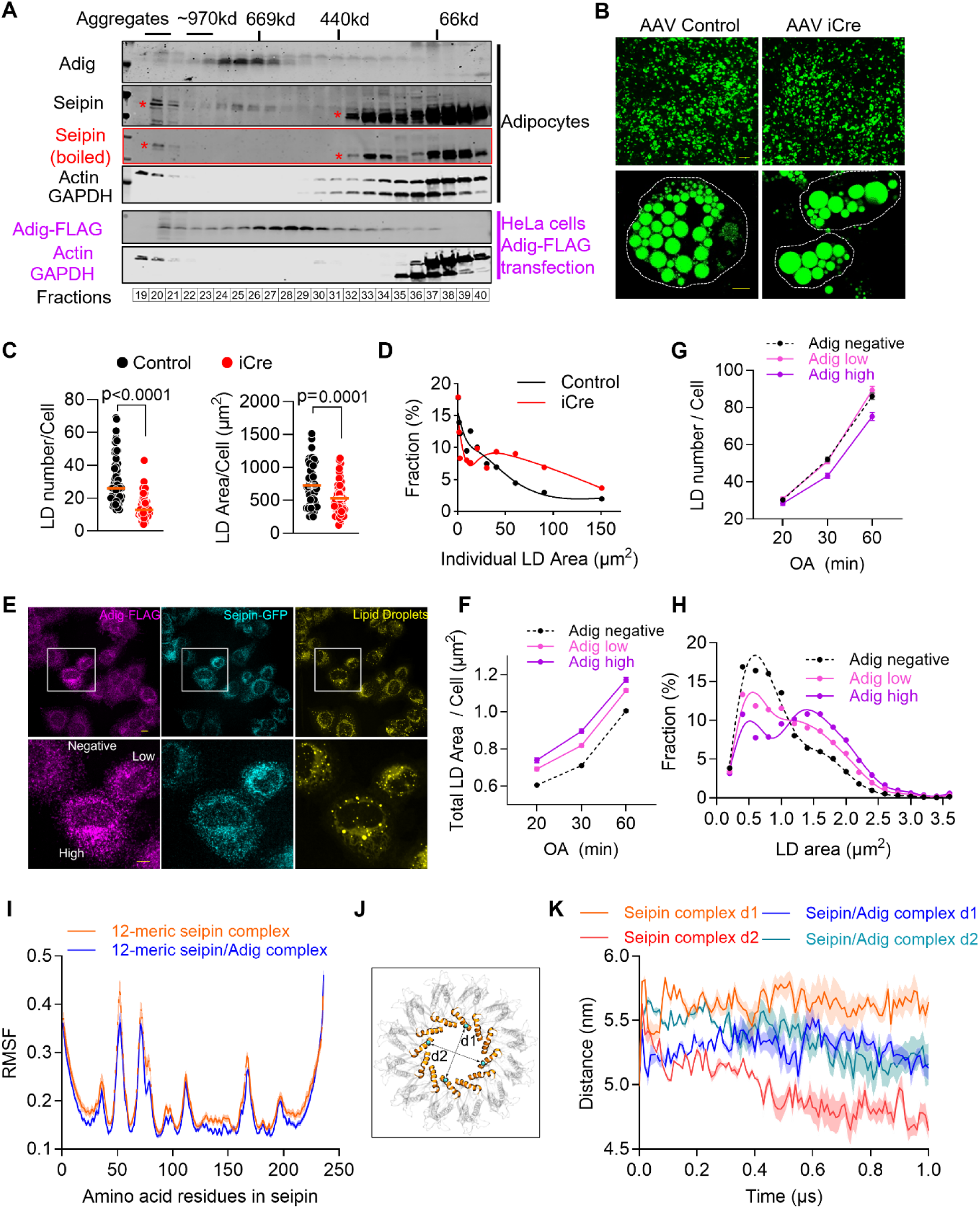
Adig facilitates the growth of lipid droplets. A. Gel filtration analysis of cell lysates from differentiated brown adipocytes and HeLa cells with Adig-FLAG overexpression. A *Superose 6 Increase* column was used to fractionate the lysates. Adig, seipin, FLAG, Actin and GAPDH antibodies were applied for Western blotting. Note the endogenous seipin band disappearing upon boiling the sample prior to analysis. The stars (*) indicate the unspecific bands. B. The elimination of Adig has no impact on the differentiation of brown adipocytes but reduces the formation of lipid droplets. SVFs were isolated from brown adipose tissue of Adig flox mice, and infected with control and iCre AAVs. Six days after adipocyte induction, cells were stained with BODIPY. Scale bar: 100µm (top), 10 µm (bottom). C. The deletion of Adig in brown adipocytes decreases the numbers and total areas of LDs in individual cells. n=63-66. An Unpaired Student’s t-test was conducted. Data are represented as mean±SEM. Only LDs more than 0.5μm^2^ were considered. D. The distribution of the LD area in brown adipocytes. A total of 1913 LDs from control cells and 983 LDs from iCre cells were counted. Only LDs more than 0.5μm^2^ were used for the calculation. Spline fitting was performed in GraphPad Prism 10.2.3. E. Adig overexpression in A431 cells promotes the growth of lipid droplets. Representative images of FLAG, GFP and LD staining after incubation with oleic acid for 30 minutes. Endogenous seipin in human epithelial A431 cells was tagged with GFP. And Adig-FLAG sable expression pool was generated. Cells were starved with lipoprotein-deficient serum (LPDS) medium overnight before oleic acid was loaded. 20, 30 and 60 minutes after treatment cells were imaged. Scale bar: 10 µm. F. Quantification of the total area of lipid droplets in each cell. Based on the expression of Adig-FLAG, cells are separated into three groups, 20 minutes: Negative (n=655), Low (n=438) and High (n=252); 30 minutes: Negative (n=698), Low (n=469) and High (n=261); 60 minutes: Negative (n=815), Low (n=536) and High (n=335). G. Quantification of the number of lipid droplets in each cell. According to the expression of Adig-FLAG, cells are separated into three groups as in (F). H. The distribution of individual LD area in A431 cells 30 minutes after oleic acid treatment. Lipid droplets from Adig negative (n=75136), low (n=78695) and high (n=61190) were represented according to individual area. I. Root mean square fluctuation (RMSF) profiles for the dodecameric Adig-free seipin and the seipin/Adig structure were calculated over 1 μs periods. The results are averaged over 3 independent simulations. The shaded area depicts the standard error of the mean (SEM). J. Schematic representation of the parameters d1 and d2 quantifying the diameter of the inner ring-like structure of the dodecameric seipin complex. d1 represents the distance between the C-alpha atom of residue 159 (shown as cyan spheres) in protomers 1 and 7; d2 represents the corresponding distance between protomers 4 and 10. The membrane-anchored helices constituting the inner ring-like structure of the dodecameric seipin complex are shown in orange, while the rest of the protein is shown as a transparent structure. K. Fluctuation of the diameters of the inner ring-like structure (measured by parameters d1 and d2) during the simulation. Results are averages of 3 independent simulations. SEM is shown as shaded area.

We assessed the function of Adig in brown and white adipocytes. To this end, we transduced brown and white adipocyte precursors isolated from Adig flox (Adig^f/f^) mice with AAV vectors expressing iCre recombinase before inducing them to undergo adipogenesis. This approach led to an efficient elimination of Adig expression in the differentiated state (**Figure S8C**). In brown adipocytes, the removal of Adig resulted in decreased expression of seipin (**Figure S8D**), but it had no effect on the adipogenic differentiation rate (**Figure 5B**). However, it is apparent that the loss of Adig decreases the total area and the total number of LDs (>0.5 µm^2^) in individual cells (**Figure 5C**). Furthermore, the percentage of supersized LDs tended to increase in Adig KO cells (**Figure 5D**). In contrast, the loss of Adig substantially impaired the adipogenesis of white adipocytes (**Figure S8E**). In the few apparent adipocytes, the total LD area and number dramatically decreased upon Adig deletion (**Figure S8F**). The size of LDs in the Adig KO cells was either very small or very large (**Figure S8G**). Combined, the deletion of Adig in adipocytes phenocopies several key observations related to LD formation made in yeast and mammalian cells in the absence of seipin.

We further tested the ability of Adig to promote LDs in human A431 cells expressing various levels of Adig. As shown in **Figure 5E**, as the expression of Adig increased, the intensity of seipin-GFP fluorescence correspondingly increased. Furthermore, the total area of LDs in individual cells was larger in overexpressing cells (**Figure 5F**). Interestingly, low Adig expression had no effect on the total number of LDs in individual cells, while high Adig expression decreased their number (**Figure 5G**). We analyzed the distribution of individual LD areas for these cells. 20, 30, and 60 minutes after exposure of cells to oleic acid (OA), Adig expression strongly increased the fraction of larger LDs (**Figure 5F**, **Figure S8H**, **and S8I**). In conclusion, the overexpression of Adig facilitates the growth of LDs by collaborating with seipin. At higher levels of Adig expression, the balance of droplet formation shifts towards fewer but larger LDs.

### Adig may support LD growth by maintaining the structural rigidity of the seipin complex

We explored in more detail the mechanisms by which Adig contributes to the seipin-mediated growth of the LD. Because Adig can promote the formation of the dodecameric but not the undecameric seipin complexes, we initially hypothesized that the inclusion of this additional protomer further enhances the efficacy in TAG sequestration. To this end, we performed coarse-grained molecular dynamics simulations of the dodecameric seipin complex embedded in a model ER bilayer enriched with 1.25 mol% TAG, as previously reported(*36*). We observed that the presence of an extra seipin protomer in the dodecameric complex did not significantly alter the efficacy of TAG sequestration within the lumen of the seipin complex when compared to the undecameric form (**Figure S8J and S8K**).

We went on to seek alternative explanations for the function of Adig in LD formation. In the Cryo-EM structure, Adig binding improved the local resolution of all segments of the dodecameric seipin complex (**Figure S5**), suggesting the rigidity of the seipin complex was increased. To test this assumption, we performed atomistic molecular dynamics simulations of seipin/Adig dodecameric complex in a 1:1 stoichiometric ratio. We observed that the root mean square fluctuation (RMSF) for the seipin/Adig structure was lower compared to the Adig-free seipin dodecameric structure, suggesting that Adig suppresses dynamic/thermal fluctuations and improves the stability of the seipin complex (**Figure 5I**). In addition, we explored the effect of the Adig association on the conformational symmetry of the inner lumen of the seipin complex by measuring the lumen diameter using two parameters, d1 and d2 (**Figure 5J**; see Methods for more details). In the seipin/Adig complex, the values for d1 and d2 converged to around 5.2 nm during the simulation, characterizing a circular symmetry for the lumen. However, without Adig, d1 and d2 varied between 4.8 and 5.6 nm, suggesting a stark deviation of the inner lumen from its regular circular symmetry (**Figure 5K**). This is supporting the view that Adig suppresses thermal fluctuations of the seipin complex.

Based on these simulations, we conclude that Adig supports LD growth more likely by maintaining the structural integrity of the seipin complex during TAG sequestration than by increasing the affinity or avidity for TAGs.

### Adig overexpression in adipocytes promotes adipose tissue expansion

Our efforts above were directed towards examining the Adig/seipin complex at the molecular and cellular levels. To probe the functional role at the level of intact tissue and for the entire organism, we resorted to gain- and loss-of-function studies in mice. We hypothesized that upregulated Adig in mouse adipose would mirror what we observed in isolated fat cells, where Adig overexpression in differentiated adipocytes can significantly increase the expression of endogenous seipin, as seen by fluorescence microscopy (**Figure S9A**). We, therefore, established a transgenic mouse model in which Adig expression can be specifically induced in adult animals in adipocytes via a Doxycycline (Dox) inducible system (Adig iTG) (**Figure S9B**). After two weeks on the Dox-containing diet, the Adig protein was significantly increased in adipose tissues (**Figure 6A**). The body weights of Adig iTG mice showed a tendency towards an increase (**Figure S9C**). Specifically, we found that the weight of brown (BAT), subcutaneous white (sWAT) and epididymal white (eWAT) adipose tissues significantly increased (**Figure 6B**). Strikingly, the LDs in BATs from Adig iTG were dramatically enlarged, as shown by H&E stains (**Figure 6C**). In WATs, the size of the adipocytes, along with the unilocular LDs, were significantly increased after Adig overexpression. (**Figure 6D and 6E**). In line with these observations, the content of triglycerides and cholesterol in BAT from iTG mice was massively elevated (**Figure 6F and 6G**). We further determined the triglyceride profiles in BAT by lipidomic analysis. Almost all the triglyceride species were increased upon Adig overexpression (**Figure 6H**), with no specific triglycerides that were disproportionally enriched compared to controls (**Figure S9D**). A triglyceride clearance test revealed that Adig overexpression significantly increased the triglyceride uptake from circulation (**Figure 6I**). We monitored the long-term effects of Adig overexpression in adipose tissues. Just like after short-term induction, we found the body weights of the Adig iTG mice increased (**Figure S9E**). Strikingly, the visible BAT from iTG mice significantly shrank (**Figure S9F**), most likely turning into white adipocytes. The residual brown adipocytes in the Adig iTG mice displayed highly-enlarged LDs (**Figure S9G**). Furthermore, the long-term Adig overexpression increased the weights of sWAT and eWAT. Both fat pads displayed cells with enlarged LDs (**Figure S9H-K**). Our results indicate that adipose tissues, especially the BAT of the Adig iTG mice, had an elevated ability to absorb triglycerides to support the growth of LDs. These observations further suggest that the enhancement of the cellular machinery for LD formation in the adipocyte is sufficient to induce beneficial effects on triglyceride metabolism systemically.

**Figure 6.**
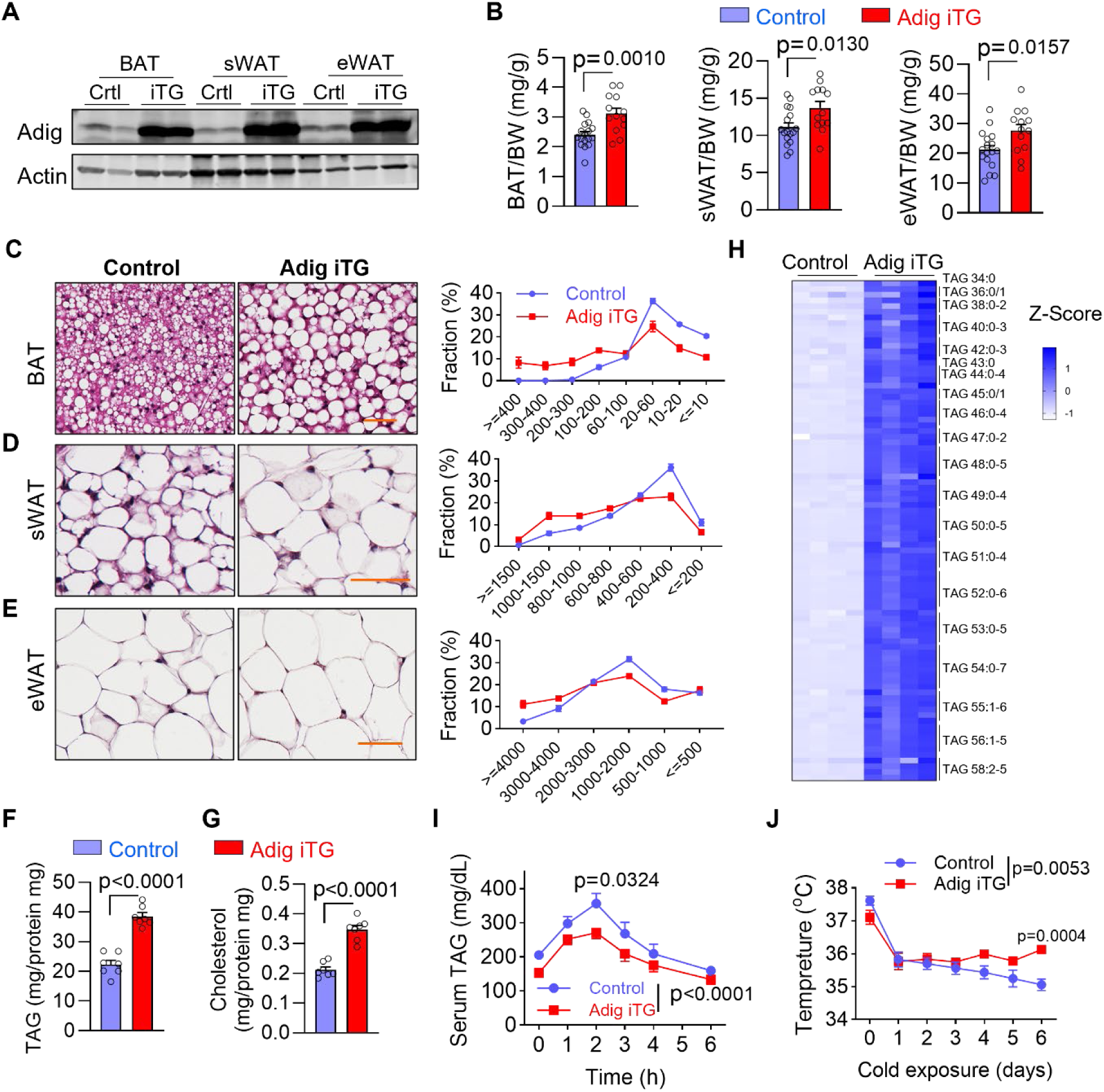
Adig overexpression promotes the expansion of adipose tissues. A. Validation for the Adig overexpression in adipose tissues. Mice were fed with doxycycline-containing chow for 2 weeks. B. Adig overexpression increases the fat mass in different depots. n=13-17. Unpaired Student’s t-test was conducted. Data are represented as mean±SEM. C. Adig overexpression increases the size of lipid droplets in brown adipose tissue (BAT). n=6. For each replicate, 1161-5600 lipid droplets were separated into 8 groups according to their size. Scale bar: 50µm. D. Adig overexpression increases the size of adipocytes in subcutaneous adipose tissue (sWAT). n=6-7. For each replicate, 214-318 adipocytes were separated into 7 groups according to their size. Scale bar: 50µm. E. Adig overexpression increases the size of adipocytes in epididymal adipose tissue (eWAT). n=7-8. For each replicate, 346-587 adipocytes were separated into 6 groups according to their size. Scale bar: 50µm. F. Adig overexpression increases the content of triacylglycerol in BAT. n=7. An Unpaired Student’s t-test was conducted. Data were represented as mean±SEM. G. Adig overexpression increases the content of cholesterol in BAT. n=7. Unpaired Student’s t-test was conducted. Data were represented as mean±SEM. H. Heat map showing lipidomic profiling for the triacylglycerols (TAGs) isolated from brown adipose tissue in control and iTG mice. The amount of TAGs was normalized to total protein, then standardized with Z score. n=4. H. Triacylglycerol (TAG) clearance test shows enhanced lipid absorption in Adig iTG mice. n=11-13. A two-way ANOVA was conducted, followed by Sidak’s multiple comparison test. Data were represented as mean±SEM. I. Adig iTG mice show enhanced thermogenesis after cold exposure. n=12. A two-way ANOVA was conducted, followed by Sidak’s multiple comparison test. Data were represented as mean±SEM.

We appreciate that in response to cold exposure, BAT drastically accelerates the uptake of triglyceride to sustain core body temperature (*37, 38*). We, therefore, tested the role of Adig during thermogenesis. We found that during 6 days of cold exposure, the drop in core body temperature in Adig iTG mice was significantly attenuated compared to control mice (**Figure 6J**), indicating Adig overexpression accelerated triglyceride uptake and boosted thermogenesis in brown adipose tissues.

### Loss of function of Adig in adipocytes impairs the growth of lipid droplets in brown adipose tissue

Two previous reports showed brown adipocyte-specific seipin deletion mice manifested aberrant LDs, cold intolerance, and impaired BAT expansion (*39, 40*). We, therefore, set out to establish a mouse model in which the Adig can be eliminated from adipocytes in a Dox-inducible manner. To this end, we first established Adig flox mice (Adig^f/f^) (**Figure S10A**) and crossed them with adiponectin-rtTA/TRE-Cre mice to generate inducible adipocyte-specific Adig knockout mice (Adig iAKO) (**Figure S10B**). To avoid the potential effects of Adig on adipogenesis, we induced the Adig KO when the mice were 5-6 weeks of age when the adipose issue was already well developed (**Figure 7A**). We examined these mice on chow diet 4 weeks after Adig deletion. The body weights of these mice were comparable to the control mice (**Figure S10C**). However, we found that most of the LDs in BATs from the iAKO mice were much smaller than the ones in Adig^f/f^ mice, although several supersized LDs were observed as well (**Figure 7B**). The accumulation of different species of triglycerides was significantly and consistently decreased in BAT from iAKO mice (**Figure 7C**). To test the impact on fatty acid uptake in different tissues, we injected ^3^H triolein into mouse arteries, and then collected blood at different time points, and finally harvested different tissues. We found iAKO mice manifested delayed clearance of ^3^H triolein (**Figure 7D**), which is in agreement with the impaired triglyceride clearance in iAKO mice (**Figure S10D**). Importantly, we found that the triglyceride uptake of BAT in iAKO mice was severely reduced (**Figure 7E**), while no significant differences were observed in the uptake into sWAT, eWAT, liver, heart and skeletal muscle (**Figure S10E-S10J**). Furthermore, when challenged with a cold exposure, Adig iAKO mice displayed an impaired gain of BAT mass compared to control mice (**Figure 7F**). Supporting the notion that Adig elimination may impair BAT function, iAKO mice displayed a lower core body temperature upon cold exposure (**Figure 7G**). In summary, the phenotype of Adig iAKO mice mimicked that of brown adipocyte-specific seipin KO mice, indicating that Adig plays a crucial role in supporting BAT function, likely through promoting seipin-driven LD formation.

**Figure 7.**
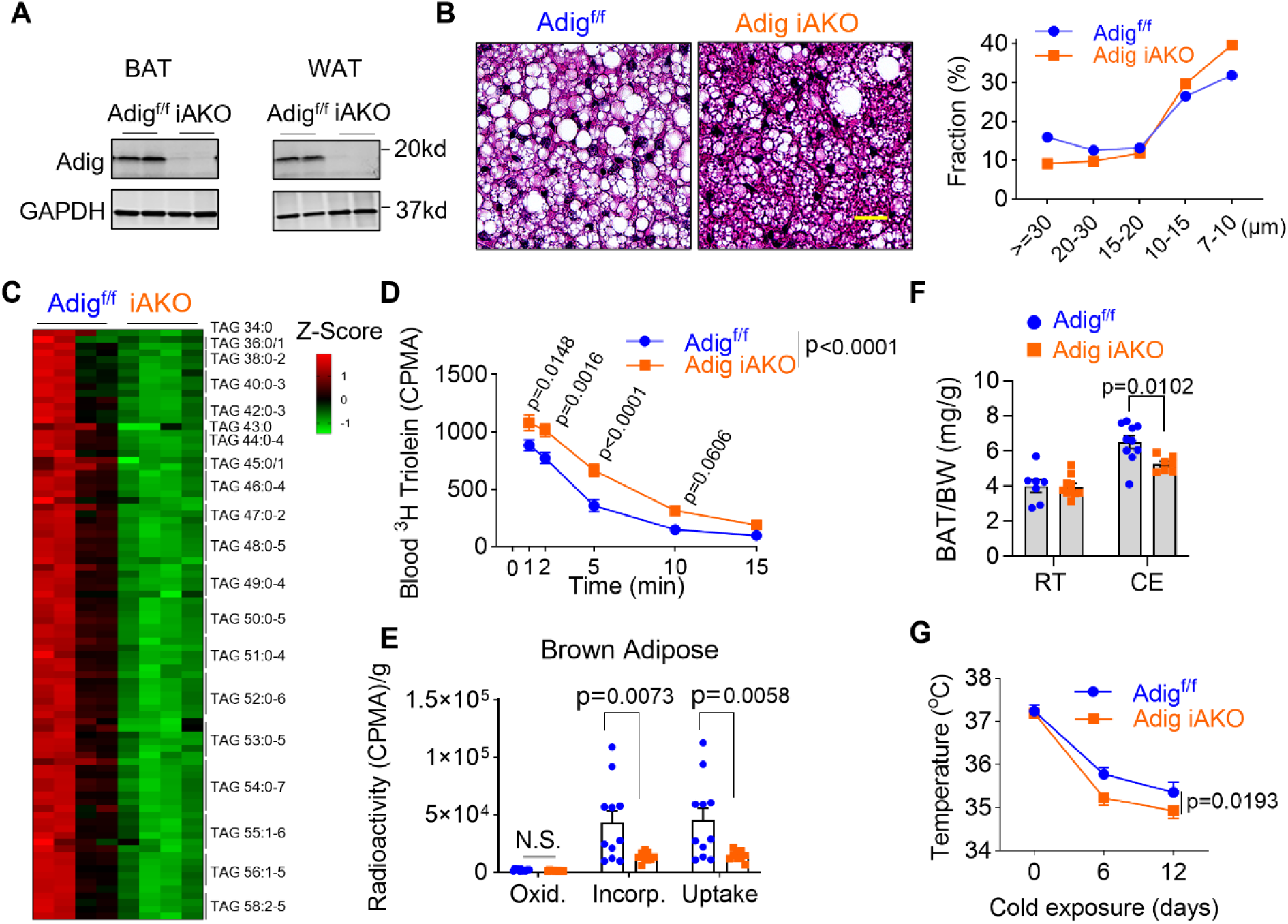
The inducible Adig elimination in adipocytes impairs the growth of lipid droplets in brown adipose tissue. A. Validation of the Adig knockout in adipose tissues. Mice were fed with doxycycline-containing chow for 4 weeks. B. Representative H.E. staining for brown adipose tissue from Adig^f/f^ and Adig iAKO mice. The elimination of Adig decreases the size of LDs in brown adipose tissue. n=4. For each replicate, 2933-10242 lipid droplets were separated into 5 groups according to their size. Scale bar, 20µm. C. Heat map showing lipidomic profiling for the triacylglycerols (TAGs) isolated from brown adipose tissue in Adig^f/f^ and iAKO mice. The amount of TAGs is normalized to the total protein amount first, then standardized with a Z score. n=4. D. Adig iAKO impairs the clearance of ^3^H Triolein from circulation. Equivalent amounts of ^3^H Triolein were injected into circulation. 1, 2, 5, 10, and 15 minutes after injection 10μl blood was collected, and the radioactive signal was measured. n=10-11. A two-way ANOVA was conducted, followed by Sidak’s multiple comparison test. Data were represented as mean±SEM. E. Adig iAKO decreases the incorporation and uptake of ^3^H-Triolein-derived fatty acids in brown adipose tissues. Oxi: oxidized fatty acids; Incorp: incorporated fatty acids; Uptake: total amount of absorbed fatty acid (oxidized and incorporated). n=9-11. A two-way ANOVA was conducted, followed by Sidak’s multiple comparison test. Data represented as mean±SEM. F. The Adig iAKO impairs the expansion of brown adipose tissues after cold exposure. n=7-10. A two-way ANOVA was conducted, followed by Sidak’s multiple comparison test. Data represented as mean±SEM. G. Adig iAKO mice display a lower body temperature during cold exposure. n=9-11. A two-way ANOVA was conducted, followed by Sidak’s multiple comparison test. Data represented as mean±SEM.

## Discussion

Here, we characterize the evolutionarily conserved microprotein Adig with regard to its subcellular localization, interaction partners, and function. We found that Adig selectively mediates the formation of previously unrecognized dodecameric seipin complex, which may reflect the actual bioactive cellular seipin complex. Furthermore, Adig stabilizes and promotes the assembly of unique dodecameric seipin complexes, therefore supporting the effective formation of LDs in cells and tissues. In mice, the expression of Adig shapes the formation of LDs and impacts the expansion and function of adipose tissues.

### Adig enhances the assembly and stability of the seipin complex

The Cryo-EM structures of the seipin/Adig complexes indicate that there are at least two segments in Adig mediating the interaction with seipin. First, the N-terminus of Adig is embedded into the wheel-like structure of the seipin complex and may interact with the previously described switch regions (*19*). Second, the α-helical stretch in Adig and the TM1 and TM2 of seipin, all of which are rich in isoleucine, leucine, and valine, intercalate into the hydrophobic Adig cluster (*41*). Based on an AlphaFold3 prediction, the key amino acid residues in Adig that putatively mediate the interactions are identified (**Figure S7**). Interestingly, these residues are evolutionarily conserved among different species (**Figure 1D**). These interactions may largely facilitate the assembly as well as enhance the stability of the overall seipin structure. Therefore, Adig expression allows more seipin foci to accumulate in cells, aiding LD formation. Additionally, Adig appears to alleviate the formation of seipin aggregates. Functionally, the stable seipin complex may be able to continually support the synthesis of triglyceride by activating DGAT2, promote the growth of LDs, and therefore enable the cells to absorb more lipids from their environment.

### Adig might be a novel component of the lipid droplet assembly complex (LDAC)

We found that endogenous Adig in adipocytes is always associated with the seipin complex, and there is strong co-localization for Adig with LDs and seipin (**Figure 5**). These observations lead us to reason that Adig might be an integral part of the lipid droplet assembly complex (LDAC) (*42*). At the same time, we have noticed that in mammalian cells without Adig, LDs can still form normally, although ectopic Adig expression can further propel LD growth. Adig elimination severely impairs the generation of LDs in adipocytes, and we speculate that Adig might form specific interactions with LDACs, especially in cells that have a high demand for lipid storage, such as adipocytes and steatotic hepatocytes. We cannot rule out that there may be an Adig homolog (yet to be identified) in other tissues that lack Adig expression.

Meanwhile, we should be cautious regarding the exact stoichiometry of seipin/Adig complex, since the Cryo-EM maps of the seipin/Adig complex represent an average based on the calculation of thousands of particles. Based on a variable co-localization of seipin and Adig in A431 cells (**Figure 4**), we cannot rule out that individual dodecameric seipin complexes associate with variable numbers of Adig components, with up to 12 Adig molecules embedded into the seipin structures, potentially regulating the activity of the overall complex.

### The dodecameric seipin complex may be a novel oligomer in mammalian cells

Seipin oligomer architecture varies in different species (*19, 28*). Two independent studies show that there are 11 copies in the human seipin complex (*20, 26*). Given the high sequence identity between human and mouse seipin, it is striking to see the undecameric and dodecameric complexes co-exist in mouse seipin. These observations raise a series of important questions. Does the endogenous form of human seipin form a dodecameric complex? Are the 11-mer or the 12-mer artifacts of overexpression in mammalian cells? If the latter is true, what is the oligomeric state of the endogenous protein? Is Adig the key factor that drives the formation of the dodecameric complex? What is the functional difference between the undecameric and dodecameric complexes? Are there any conformational changes seen in the dodecameric seipin complex during LD growth as proposed in the yeast decameric seipin structure (*19*)?

Although resolving these questions is beyond the scope of our current study, our results provide some valuable insights. Adig can only be found in the dodecameric seipin complex (**Figure 3**). In human A431 cells, Adig can stabilize and interact with endogenous human seipin (**Figure 4**). Taken together, these results indicate that the dodecameric seipin complex likely exists in its native conformation in human cells. From the perspective of structural biology and predictions, compared to undecamer, the dodecameric form has more hydrophobic TMs -and luminal hydrophobic α-helix residues and has more space in the middle of the cage-like structure, and Adig binding features a more rigid seipin complex, all of which may increase its capacity to concentrate and store triglycerides. In support of this prediction, LDs grow faster and bigger with Adig overexpression in A431 cells (**Figure 5**). Our results support the notion that the novel dodecameric seipin complex identified here is a functionally important oligomer in mammalian cells.

### Seipin TM segments are visible in the Cryo-EM structure of seipin/Adig complex

Previous molecular simulation studies revealed that the TM segments in human seipin play a key role in attracting triglycerides, prompting LD budding (*36, 43, 44*). However, due to the lack of details on the atomic structure of the TM segments in previous reports, there were concerns about the accuracy of these simulations. Recently, two studies of the yeast seipin orthologue Sei1 successfully resolved the TM structure (*19, 28*). Importantly, one study proposed a conformation switch model in TMs to explain the process of LD growth and budding (*19*). However, given the sequence diversity, the yeast TM structure may not reflect the structure of human seipin. Moreover, the conformational switch model is incompatible with the current human undecameric seipin complex. Our study may pave the way to solve these issues: First, with the addition of Adig to the seipin complex, the TM segments are stabilized and, therefore, captured in Cryo-EM, which gives us the opportunity to add TM segments to molecular simulations. Second, the seipin/Adig Cryo-EM structure may be able to be captured at different stages of LD formation, which can test whether the TMs in our dodecameric structure follow a similar mechanism as in the yeast 10-meric seipin. Third, the stabilized seipin structure may allow the resolution of the entire LDAC. Currently, we still do not have any insights as to how LDAF1 works inside the seipin oligomers (*26*), maybe because previous efforts may have relied on the undecameric seipin structure with considerable flexibility of the TMs. Here, we found the seipin/Adig 12-mer complex is relatively rigid, and it is easier to resolve its structure at a higher resolution. Therefore, we might have a better chance to resolve the seipin/Adig/LDAF1 triple complex structure in the future.

### Physiological and pathological implications for the function of Adig in Seipin complex

Depending on the tissue context, seipin may display different functions. In humans, the seipin null mutant is responsible for the most severe form of congenital generalized lipodystrophy, highlighting its vital role in adipogenesis and LD formation (*31*). In the testis, seipin deletion causes teratozoospermia (*45*). Seipin is highly expressed in the human nervous system. Several missense mutants around the glycosylation sites lead to aberrant seipin aggregation, impair motor neurons and lead to neurological seipinopathy manifesting in muscle weakness, deafness, and dementia (*32, 46*). In brown adipose tissue, seipin is absolutely essential for the expansion of brown adipocytes (*39, 40*). It is still unknown whether all these abnormalities are related to defects in LD formation. Probably, other mechanisms, including phospholipid metabolism (*47*) and/or Ca^2+^ homeostasis (*48, 49*), are involved as well. Nevertheless, it is essential to keep seipin properly folded, oligomerized, and correctly located under all of the conditions. In our study, we identify the microprotein Adig as an important “chaperone-like” regulator for the seipin complex. We propose that through modulating the expression of Adig, the activity of the seipin complex can be accurately regulated, which presents a potential opportunity to treat seipin-related diseases.

## Acknowledgements

We thank Drs. Thomas Gillette, Shiqian Li, Xiaofeng Qi, and Abel Szkalisity for comments and suggestions. For support with Cryo-EM studies, we thank the Structural Biology Lab and Cryo-EM Core Facility at UT Southwestern Medical Center, which are supported by grant RP220582 from the Cancer Prevention & Research Institute of Texas (CPRIT). We thank the Electron Microscopy Core Facility at UTSW for the assistance in Apex2 staining and Biomedicum Imaging Unit, supported by HiLIFE and Biocenter Finland, for support in light microscopy. We thank Proteomic Core at UTSW for the identification of Adig interactive proteins. We thank the mouse Transgenic Core at UTSW for the generation of TRE-Adig and Adig flox mice. We thank the Metabolic Phenotyping Core at UTSW for the lipidomic study and the measurements of triglycerides and cholesterol. We thank CSC - IT Center for Science Ltd (Espoo, Finland) for providing computing resources to perform molecular dynamics simulations.

## Author contributions

Conceptualization: CL, PES

Methodology: CL, XNS, JBF, LV, WK, YangL, YH, EI, ChenL

Investigation: CL, XSN, JBF, LV, NJ, YanL, XP, MP, CJ, RG, CV, LS, SC, JV, AC, DLP, MYW, TO, ARN, RMW

Visualization: CL, XNS, LV, YanL, XP, MP, OV

Funding acquisition: PES, EI, IV, DYO

Project administration: PES, JMG, YH

Supervision: PES, EI, ZVW, JMG

Writing – original draft: CL

Writing – review & editing: CL, JBF, DYO, SZ, ZVW, JMG, RMW, YH, PES

## Supplementary Materials

Materials and Methods

Figure S1 to S10

Tables S1 and S2

---

**Figure S1.**
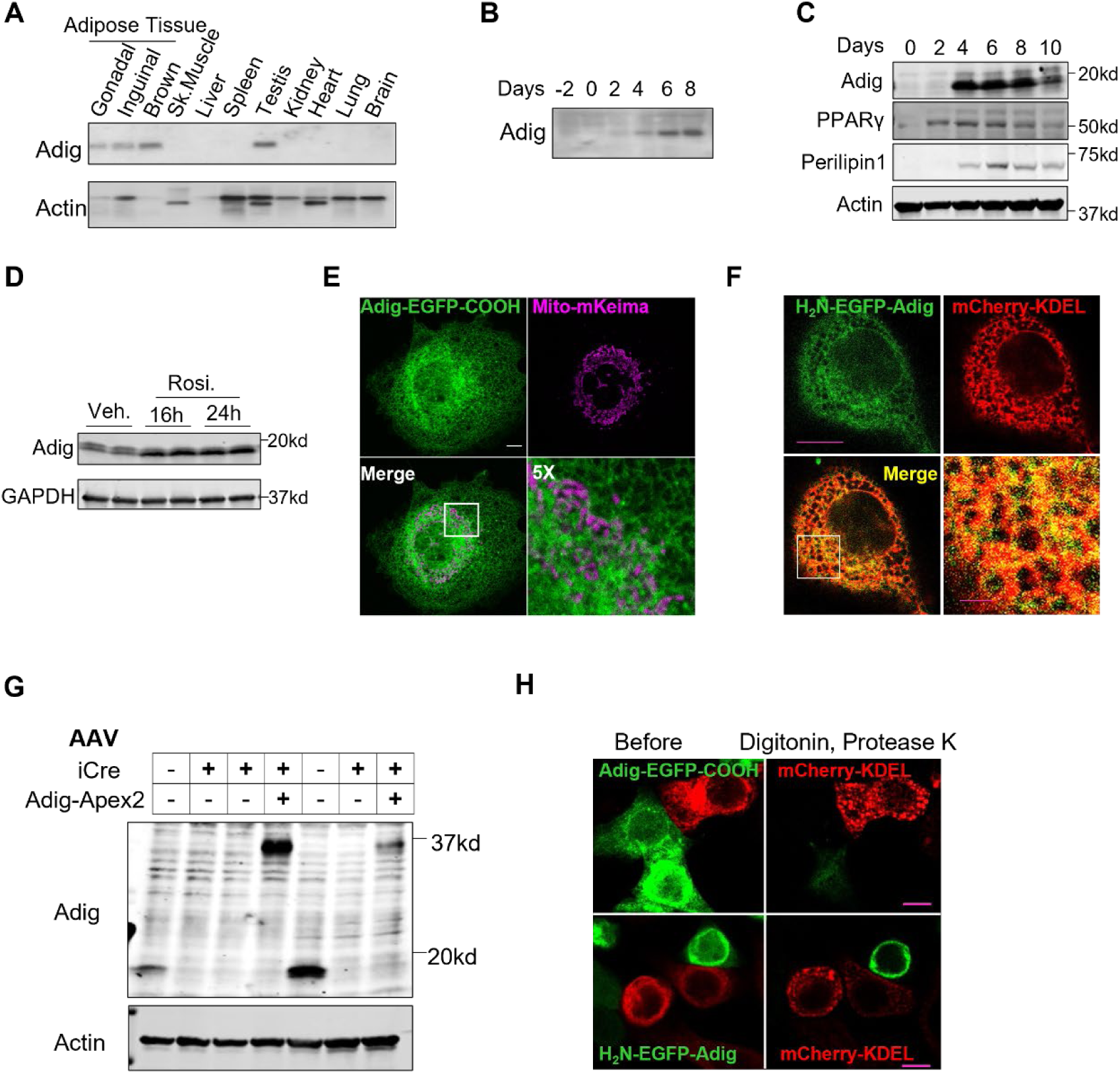
Adipogenin is highly expressed in adipocytes and localized to the ER. A. Northern blot analysis for the transcriptional profile of Adipogenin (Adig) in different tissues. Note: Adig mRNA is highly expressed in Adipose tissues and testis. B. Representative Northern blot shows induction of Adig during differentiation of 3T3-L1 from fibroblasts to adipocytes. C. Adig protein is strongly upregulated during adipogenesis in white adipocytes. D. Rosiglitazone (Rosi) elevates the protein expression of Adig in brown adipocytes. Nine days after induction, Rosi was added into medium to treat cells for 16 and 24 hours. E. Adig is not localized to mitochondria. Protein mito-mKeima indicates mitochondrial localization in COS-7 cells. Scale bar: 10µm. F. Adig is co-localized with ER marker KDEL. Plasmids EGFP(N)-Adig and ER-mCherry-KDEL are transfected into HeLa cells. The picture was taken from cells with low expression of EGFP(N)-Adig. Scale bar: 10µm. G. Assessment of the expression in Adig and Adig-Apex2 in adipocytes infected with AAV-EGFP, AAV-iCre and/or AAV-Adig-Apex2. H. Representative images show the Fluorescence Protease Protection (FPP) Assay. Scale bar: 2µm.

**Figure S2.**
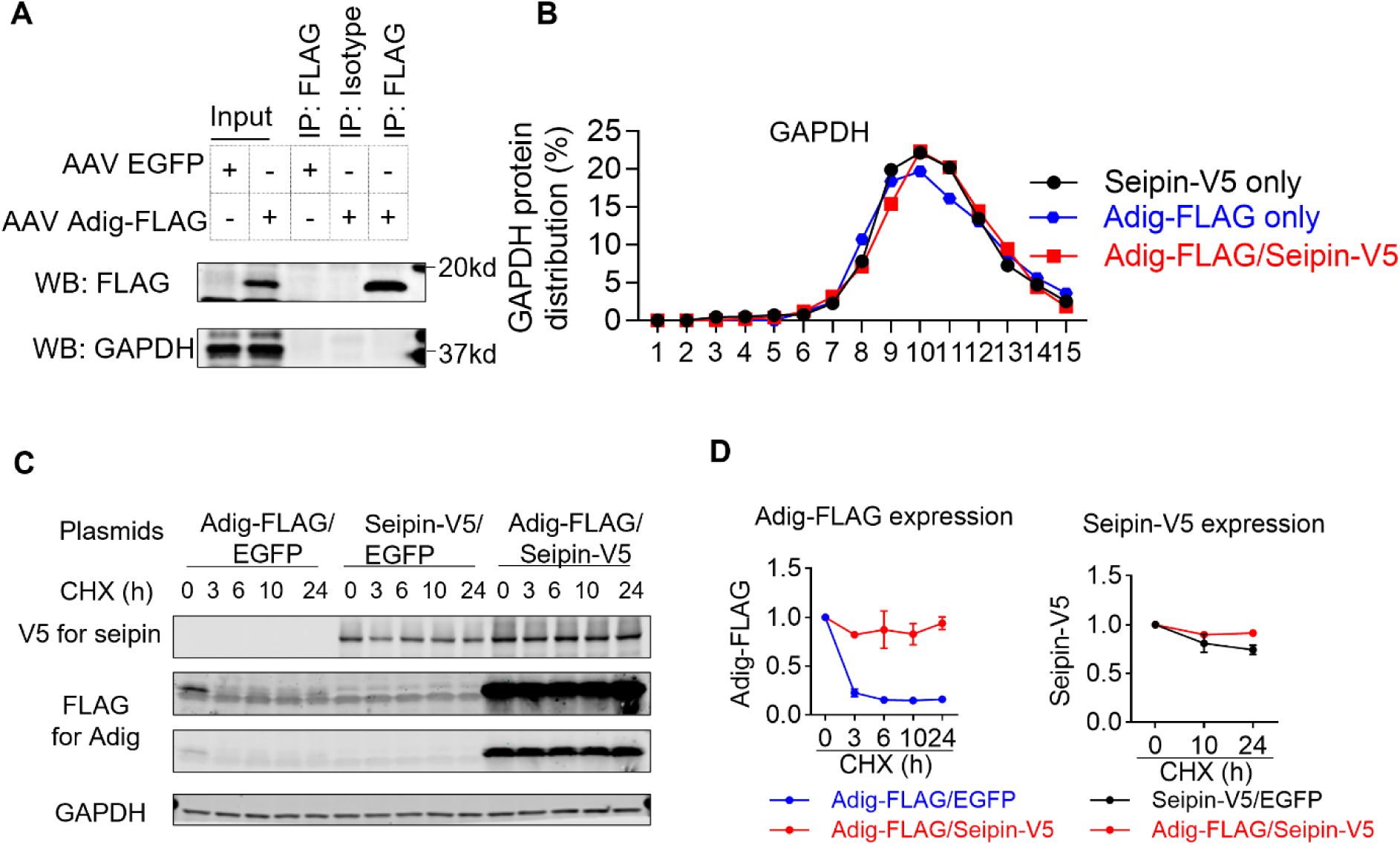
Adig interacts with and stabilizes seipin. A. Validation for FLAG immunoprecipitations in adipocytes. B. Distribution analysis for the expression of GAPDH in fractions from Figure 2C-2E. The expression of GAPDH was quantified by LI-COR Image Studio Lite first, and the percentage was calculated by dividing the individual expression in each fraction with the summing-up expression from all fractions. C. Western blot analysis for the degradation of Adig-FLAG and Seipin-V5. Different combinations of plasmids are transfected into HEK293T cells. Two days later, cycloheximide (CHX) was added into medium. Cell lysates were harvested at the indicated time points. D. Quantification for the expression Adig-FLAG and Seipin-V5 from (C). In each group the individual expression of Adig or V5 was normalized to the level at the first time point.

**Figure S3.**
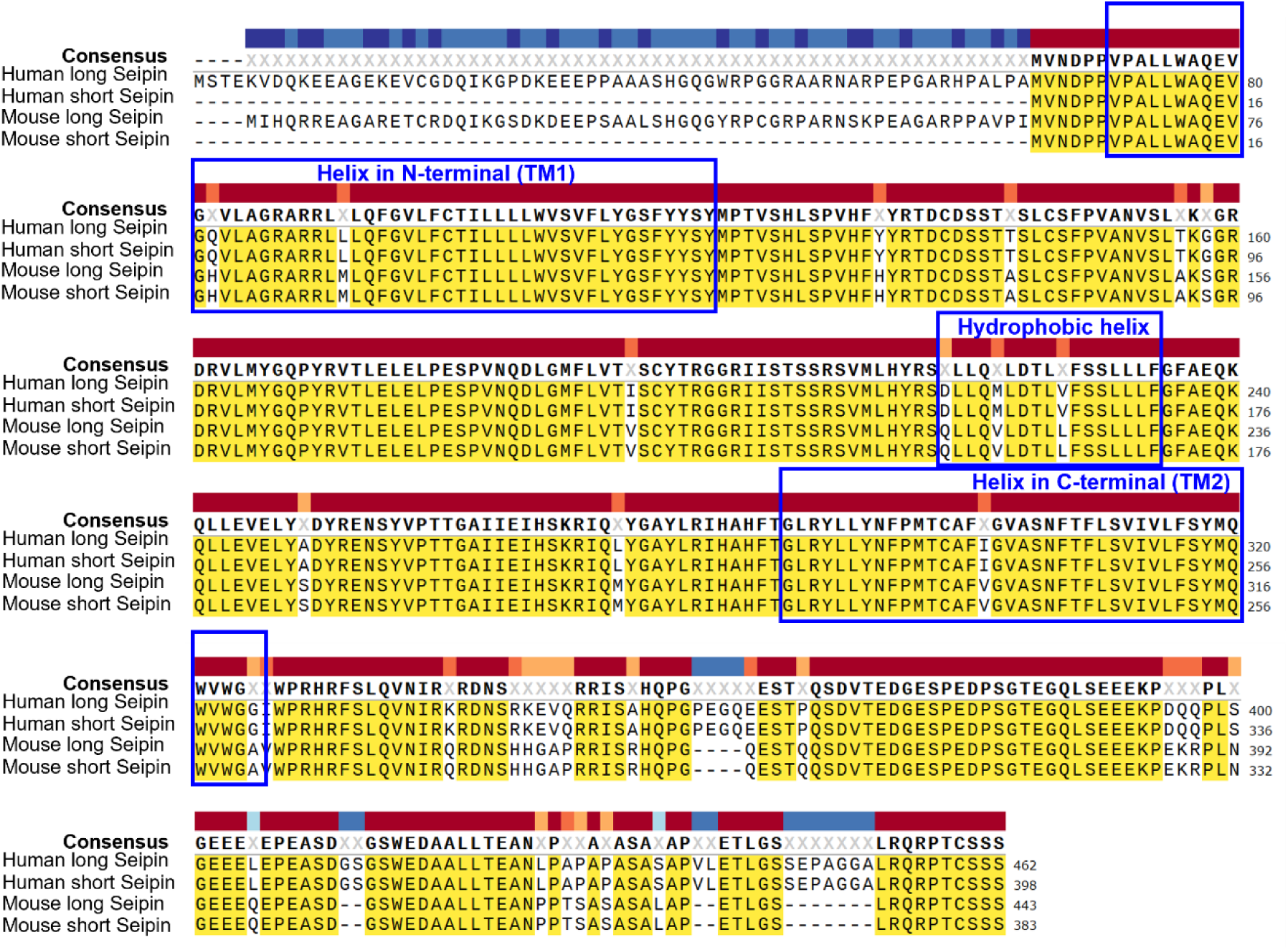
Alignment of amino acid sequences of human and mouse seipin isoforms. Mouse and human Seipin show high conservation in amino acid sequences. The N-terminal transmembrane segment (TM1), the luminal hydrophobic helix and the C-terminal transmembrane segment (TM2) are highlighted.

**Figure S4.**
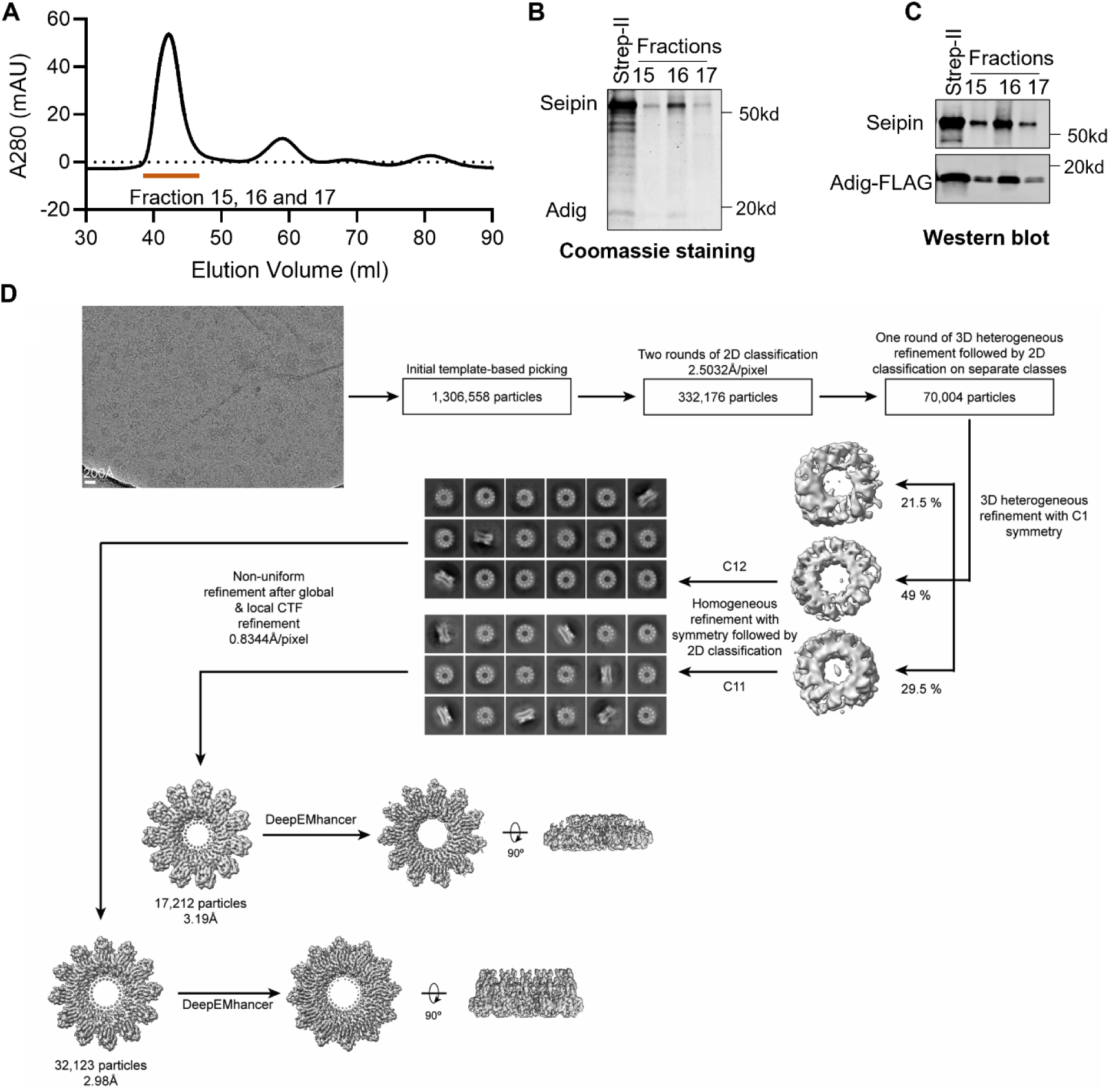
Cryo-EM structure of mouse seipin/Adig complex. A. The seipin/Adig complex was purified from the Expi293 cells as described in the methods. After strep-II affinity purification, the complex was applied to size-exclusion chromatography. Fractions 15-17 showed a sharp peak in UV absorption. B. Coomassie staining for the products from the strep-II affinity purification and fraction 15-17 in (A). The Seipin and Adig bands were identified by their respective molecular weight. C. Western blot analysis for the products from the strep-II affinity purification and fraction 15-17. Seipin and Adig were visualized by seipin and FLAG antibodies. D. Schematic showing Cryo-EM data processing steps for the seipin/Adig complex. A representative micrograph is shown.

**Figure S5.**
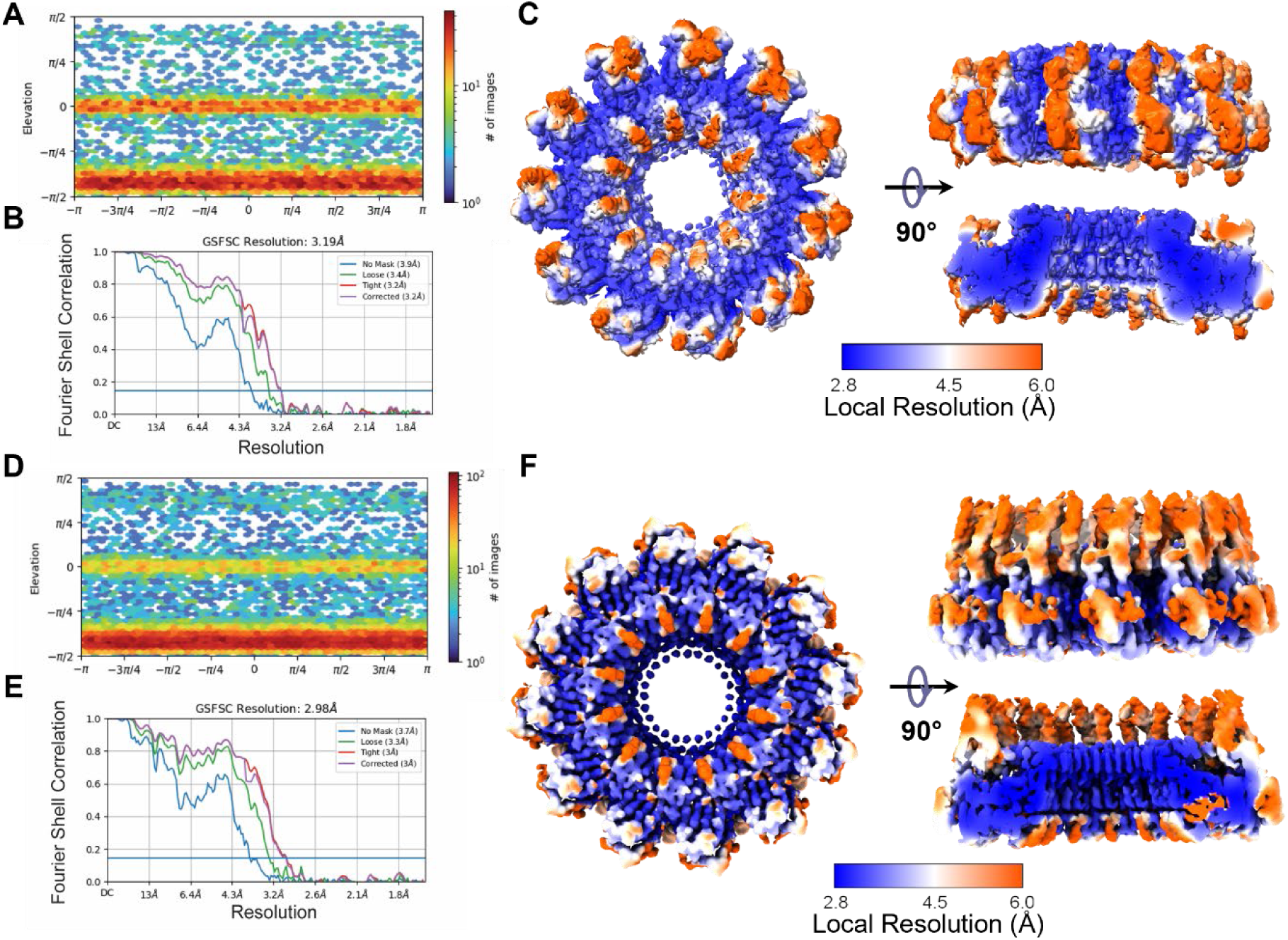
Resolution estimation for the seipin undecameric complex and seipin/Adig dodecameric complex. A. Euler angle distribution plots for the seipin undecameric complex. B. Fourier Shell Correlation (FSC) plots for the seipin undecameric complex. C. Local resolution maps of the seipin undecameric complex. D. Euler angle distribution plots for the seipin/Adig dodecameric complex. E. Fourier Shell Correlation (FSC) plots for the seipin/Adig dodecameric complex. F. Local resolution maps of the seipin/Adig dodecameric complex.

**Figure S6.**
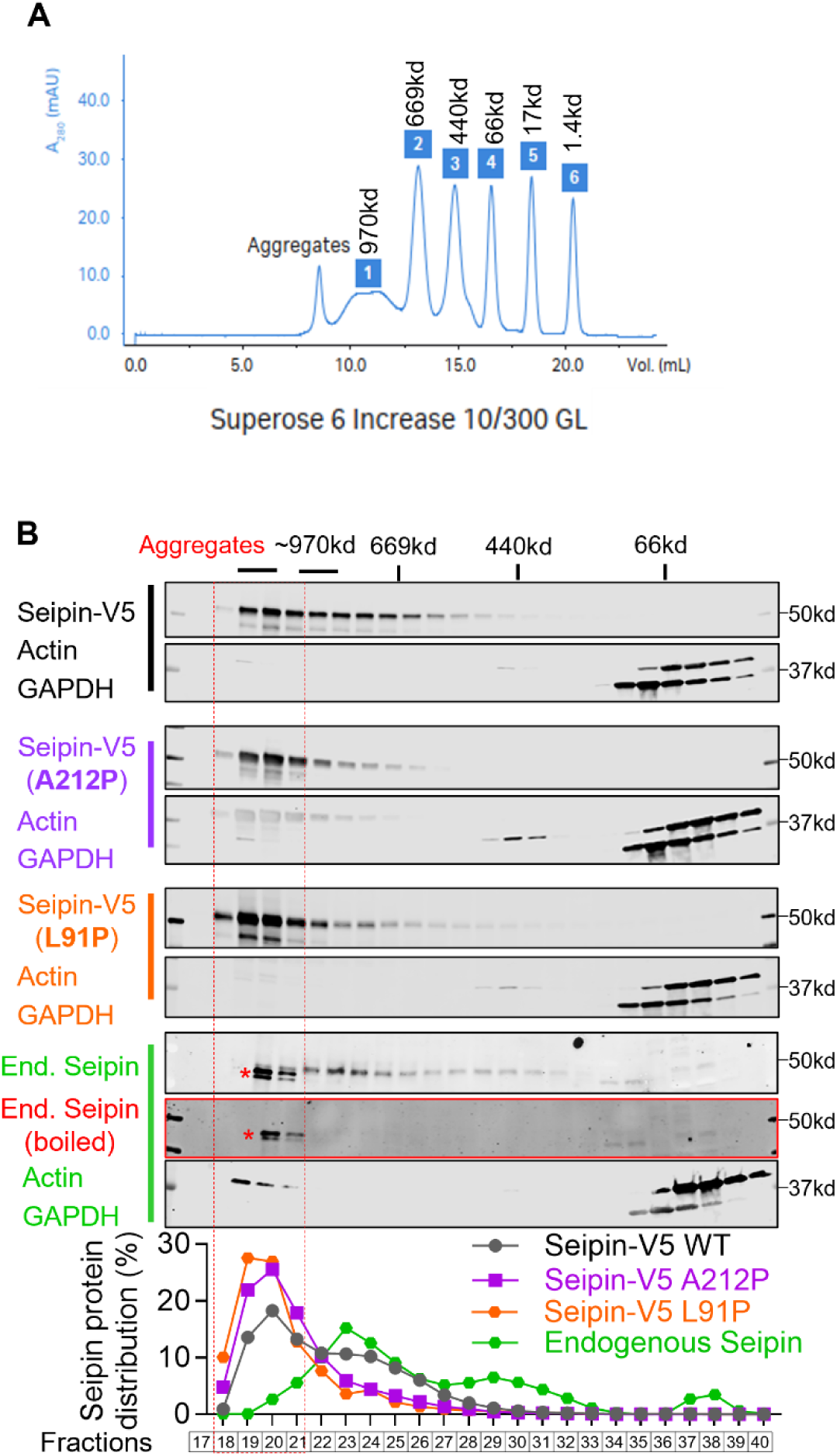
Adig promotes the assembly of the seipin complex. A. The distribution of 6 standard proteins in size-exclusion chromatography with the *Superose 6 Increase 10/300 GL* column. Data from the Cytiva^TM^ official data file. B. Gel filtration analysis for cell lysates from HeLa cells and HEK293T cells with Seipin-V5, Seipin-V5 (A212P) and Seipin-V5 (L91P) overexpression. To eliminate the seipin band in Western blots, samples were boiled at 98 C° for 10 minutes. The stars (*) indicate the nonspecific bands. A *Superose 6 Increase* column was used to fractionate the elution. V5, seipin, Actin and GAPDH antibodies were applied for Western blotting. The expression of V5 and seipin were quantified by LI-COR Image Studio Lite first, and the percentage was calculated by dividing the individual expression in each fraction with the sum of all fractions. Note the mutated seipin accumulated more in the fractions containing aggregates.

**Figure S7.**
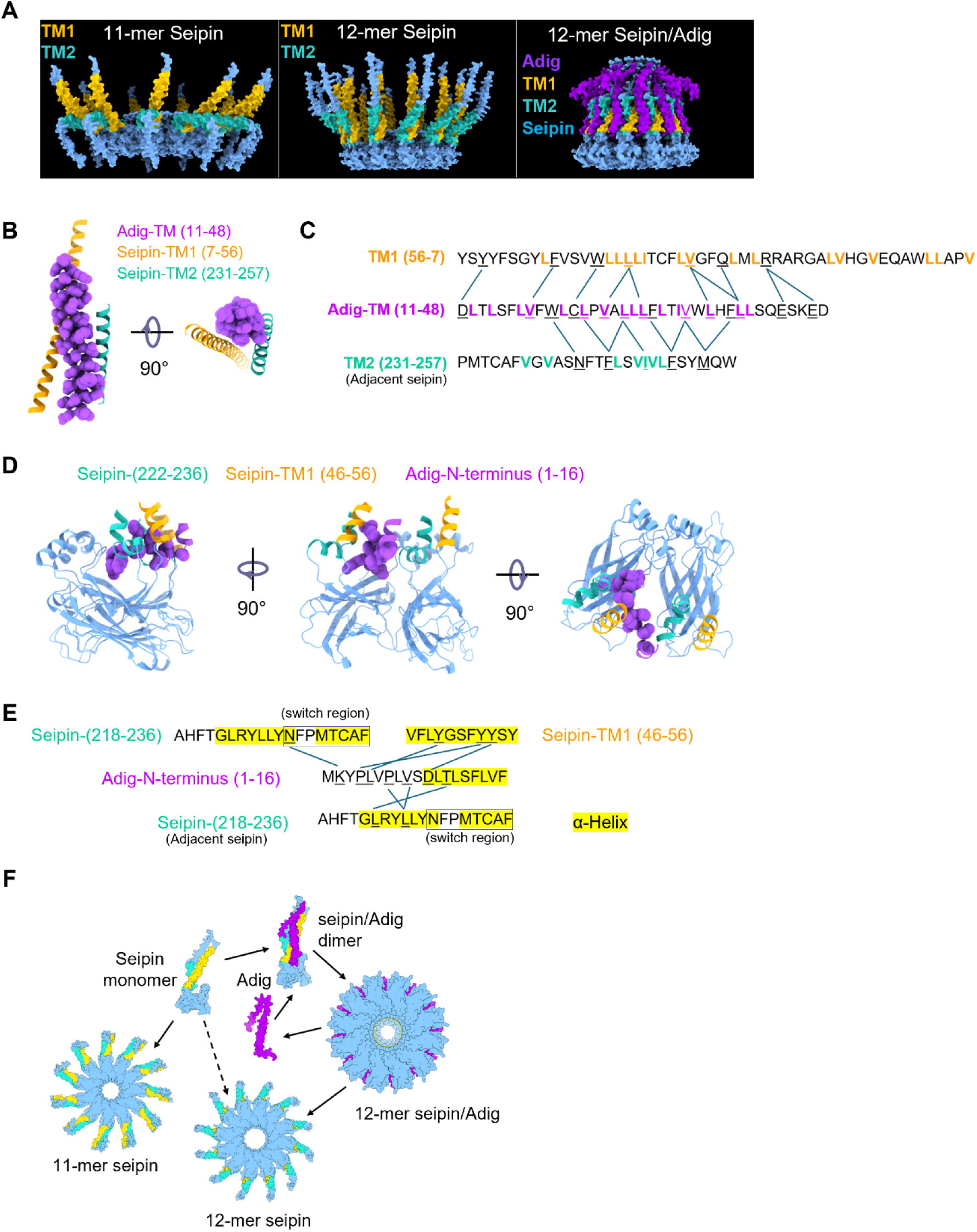
The Seipin/Adig complex modelled by AlphaFold 3. A. AlphaFold 3 predictions for the structures of undecameric seipin complex, the dodecameric seipin complex and the dodecameric seipin/Adig complex. The complete sequence of mouse Adig and most of the sequence of mouse seipin (1-282, short isoform) were uploaded into AlphaFold 3. The N-terminal transmembrane segment (TM1) and the C-terminal transmembrane segment (TM2) in seipin are colored brown and cyan respectively. Adig is colored purple. B. The TM of Adig clustered with the TM1 and the TM2 from two adjacent seipin copies. This structure is based on the predictions generated by AlphaFold 3. TM1 and TM2 in seipin are colored brown and cyan respectively. The TM in Adig is colored purple. The side chains in Adig are shown. C. Putative interaction network between the transmembrane segments from Adig and seipin. The potential interaction between residues is deduced from their proximity. D. Adig N-terminus interacts with the curvature regions of TM2 from two adjacent seipin copies. The side chains in Adig are shown. E. Putative interaction network between N-terminus of Adig with the switch region, TM1 and TM2. The potential interaction between residues is deduced from their proximity. F. Putative model for the process of seipin assembly. Seipin and Adig monomers may bind each other following their individual translation. The resulting seipin/Adig heterodimers may further fold into 12-mers. Seipin monomers may also directly assemble into 11-mers or 12-mers.

**Figure S8.**
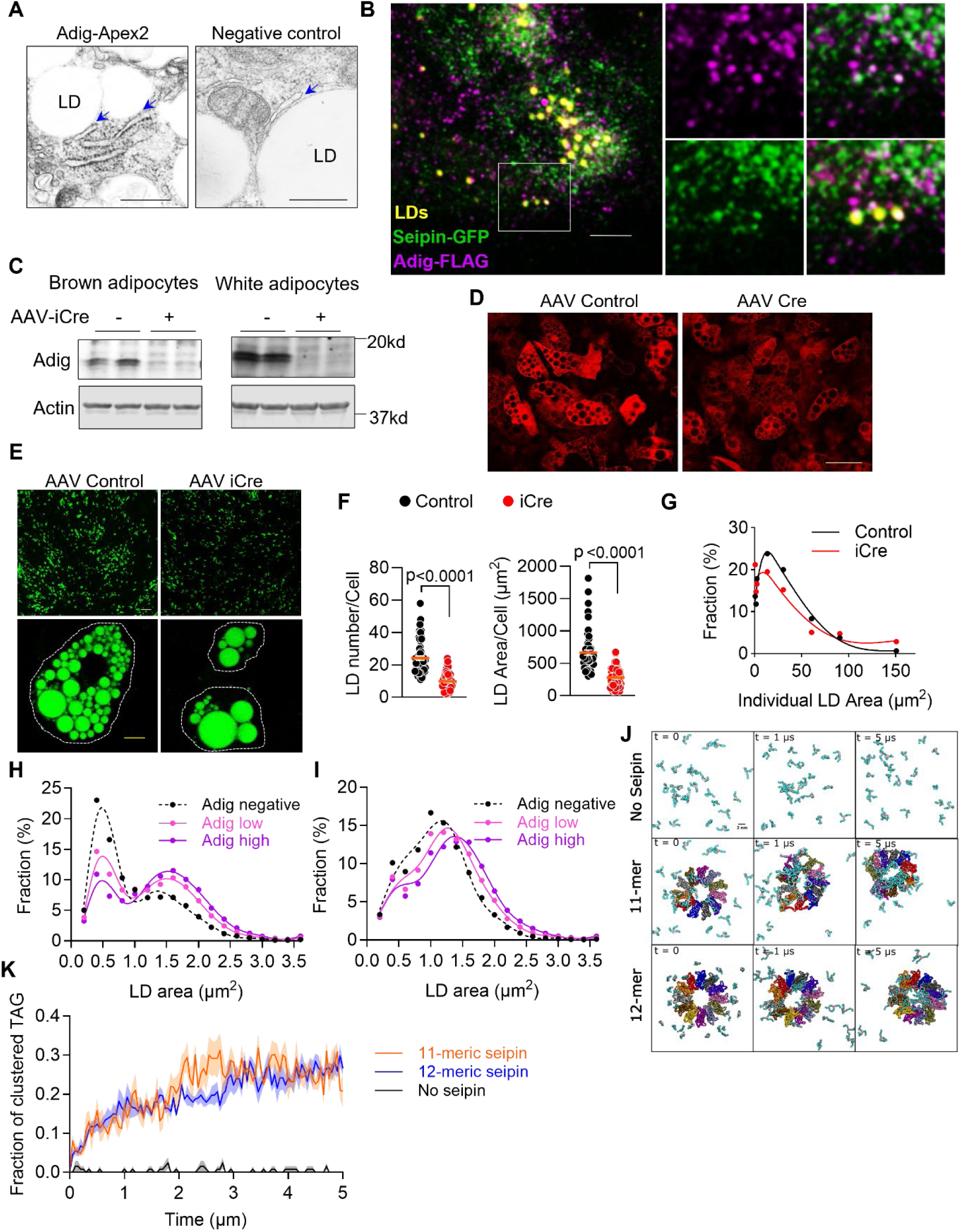
Adig facilitates the growth of lipid droplets. A. Adig-Apex2 staining in adipocyte. Stromal vascular fractions (SVFs) were isolated from adipose tissue in Adig flox mice and then differentiated into mature adipocytes. To make sure Adig-Apex2 was expressed at physiological levels, preadipocytes were infected with AAV-iCre and AAV-Adig-Apex2. For the negative control group, preadipocytes were infected with AAV-EGFP or AAV-iCre. Scale bar: 500nm. Blue arrows indicate Apex2 signals in the contacts between ER and lipid droplet. B. The co-localization of seipin-GFP, Adig-FLAG and LDs. Endogenous seipin in human epithelial A431 cells was tagged with GFP, and an Adig-FLAG stably expressing pool was generated. Cells were starved with lipoprotein-deficient serum (LPDS) medium overnight before oleic acid loading for 10 minutes. Scale bar: 10μm. C. Validation for the deletion of Adig in adipocytes. SVFs were isolated from adipose tissue in Adig flox mice and were infected with or without AAV-iCre before induction. D. Immunofluorescence of endogenous seipin in brown adipocytes with or without AAV iCre. Scale bar: 50μm. E. Adig KO inhibits the adipogenesis of white adipocytes and reduces the formation of lipid droplets. SVFs were isolated from subcutaneous adipose tissue in Adig flox mice and infected with control and iCre AAVs. Six days after adipocyte induction, cells were stained with BODIPY. Scale bar: 100µm (top), 10 µm (bottom). F. Adig deletion in white adipocytes decreases the numbers and total areas of LDs in individual cells. n=59-60. Only LDs more than 0.5μm^2^ were calculated. An Unpaired Student’s t-test was conducted. Data are represented as mean±SEM. G. The distribution of the LD areas in white adipocytes. A total of 1427 LDs from control cells and 590 LDs from iCre cells were counted. Only LDs more than 0.5μm^2^ were considered. Fit spline is performed in GraphPad Prism 10.2.3. H. The distribution of individual LD areas in A431 cells 20 minutes after oleic acid treatment. Lipid droplets from Adig negative (n=38261), low (n=54368) and high (n=52167) were fractioned according to individual area. I. The distribution of individual LD areas in A431 cells 60 minutes after oleic acid treatment. Lipid droplets from Adig negative (n=120470), low (n=116767) and high (n=76793) were fractioned according to individual areas. J. Representative snapshots (top view) of coarse-grained simulations with 1.25 mol% TAG in the membrane bilayer at different time points (t) during the simulations. Acyl chains of TAGs are shown in cyan; glycerol moiety is depicted in light pink. Each protomer of the seipin oligomer is shown with a different color. Other membrane lipids, water, and ions are not shown for clarity. K. Analysis of the simulations shown in (J), depicting the fraction of TAGs clustered together. The values are averages of 5 independent simulations. Data are represented as mean±SEM. SEM is shown as shaded region.

**Figure S9.**
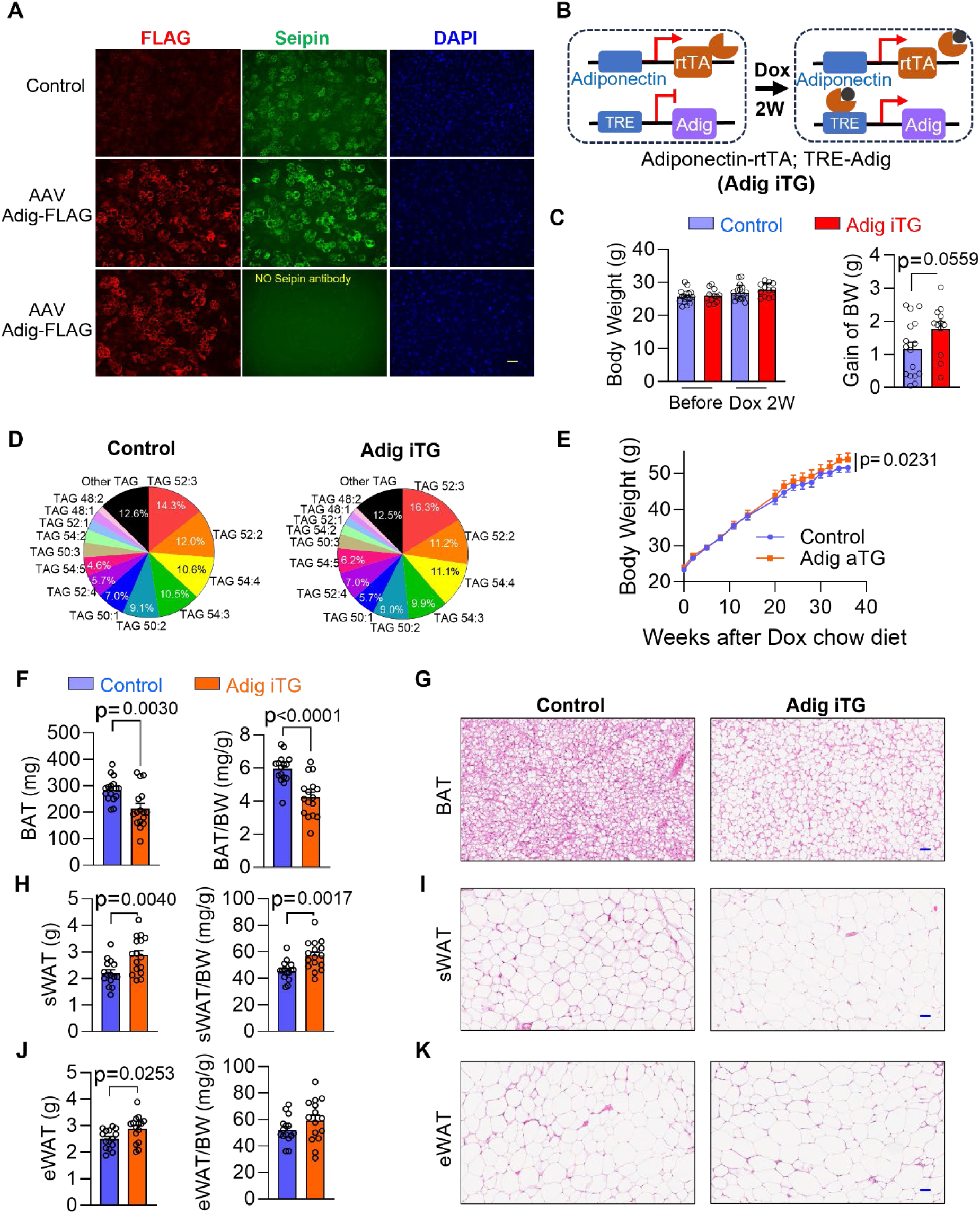
Adig overexpression promotes the expansion of adipose tissues. A. Immunofluorescence of endogenous seipin in adipocytes infected with or without AAV Adig-FLAG. Scale bar, 50μm. B. Scheme showing the generation of inducible Adig overexpression mice. Adiponectin-rtTA and TRE-Adig double-positive mice (Adig iTG) were fed with doxycycline-containing chow for 2 weeks. Adiponectin-rtTA or TRE-Adig single-positive mice were used as controls. C. The change of body weight before and after doxycycline (Dox)-containing chow for 2 weeks. n=12-17. An Unpaired Student’s t-test was conducted. Data are represented as mean±SEM. D. The distribution of the different triglyceride species in in the lipidomic analysis displayed in Figure 6H. E. The body weight gain during long-term expression of Adig in adipose tissues from control and Adig iTG mice. n=12-14. A two-way ANOVA was conducted, followed by Sidak’s multiple comparison test. Data represented as mean±SEM. F. Long-term Adig overexpression leads to shrinkage of BAT mass. n=16. An Unpaired Student’s t-test was conducted. Data represented as mean±SEM. G. Representative H&E staining for the BAT from control and Adig iTG mice. Scale bar, 50μm. H. Long-term Adig overexpression increases the mass of sWAT. n=15-16. An Unpaired Student’s t-test was conducted. Data represented as mean±SEM. I. Representative H&E staining for the sWAT from control and Adig iTG mice. Scale bar, 50μm. J. Long-term Adig overexpression increases the mass of eWAT. n=15. An Unpaired Student’s t-test was conducted. Data are represented as mean±SEM. K. Representative H&E staining for the eWAT from control and Adig iTG mice. Scale bar, 50μm.

**Figure S10.**
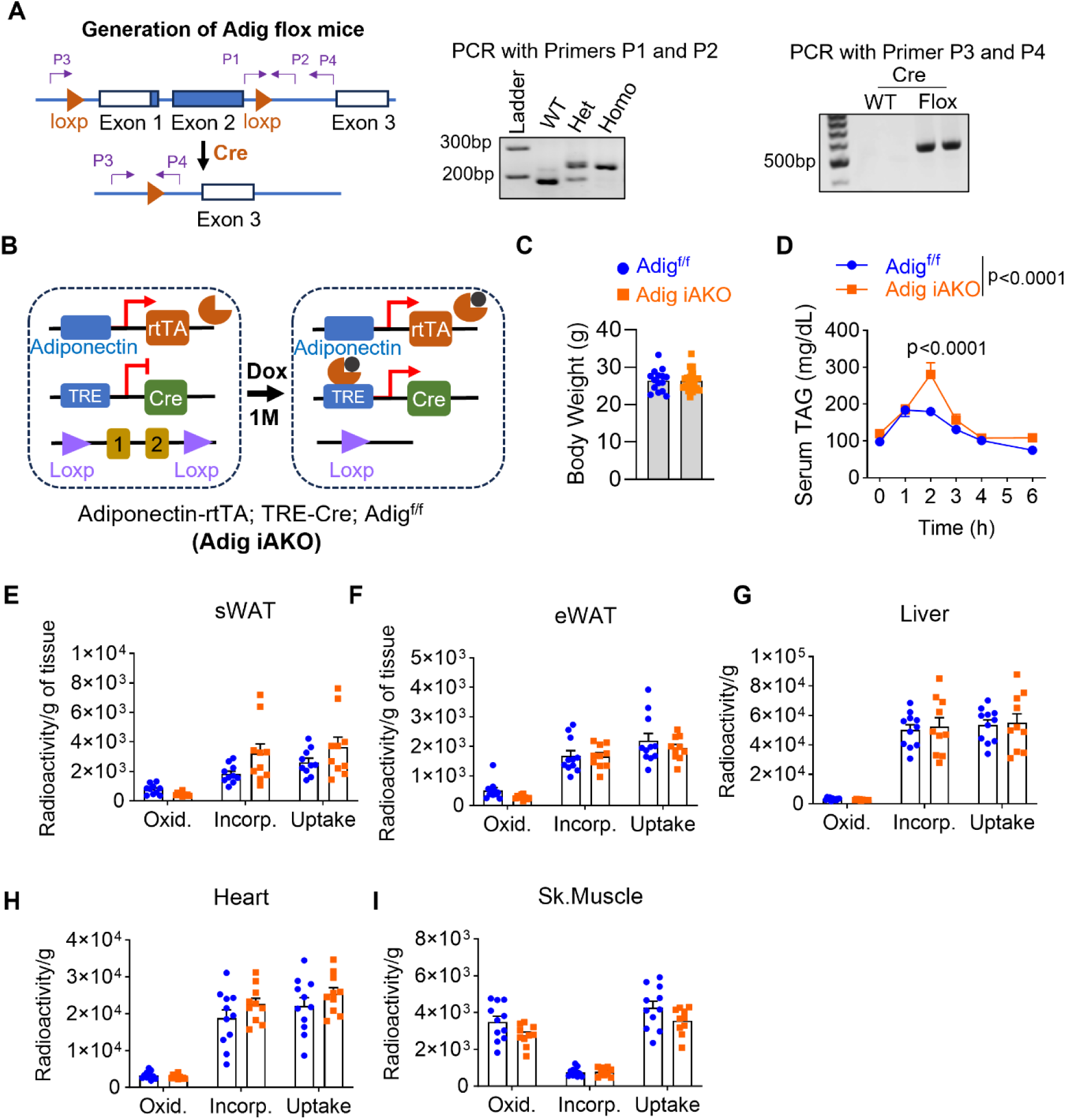
An inducible Adig knockout in mature adipocytes impairs the growth of lipid droplets in brown adipose tissue. A. Scheme showing the generation of Adig flox mice. B. Scheme showing the generation of inducible Adig knockouts in mature adipocytes. Adiponectin-rtTA and TRE-Cre double-positive mice were fed with doxycycline (Dox)-containing chow for 4 weeks. Adiponectin-rtTA positive and TRE-Cre negative mice were used as controls in the Adig^f/f^ mice. C. The body weighs of Adig^f/f^ and Adig iAKO mice fed with Dox containing chow for 4 weeks. D. A triacylglycerol (TAG) clearance test shows impaired lipid absorption in Adig iAKO mice. n=10-12. A two-way ANOVA was conducted, followed by Sidak’s multiple comparison test. Data represented as mean±SEM. L. The uptake of ^3^H Triolein-derived fatty acids in subcutaneous white adipose tissue (sWAT) from Adig^f/f^ and iAKO mice. n=10. M. The uptake of ^3^H Triolein-derived fatty acids in epidydimal white adipose tissue (eWAT) from Adig^f/f^ and iAKO mice. n=10-11. N. The uptake of ^3^H Triolein-derived fatty acids in liver from Adig^f/f^ and iAKO mice. n=10-11. O. The uptake of ^3^H Triolein-derived fatty acids in heart from Adig^f/f^ and iAKO mice. n=10-11. P. The uptake of ^3^H Triolein-derived fatty acids in skeleton muscle from Adig^f/f^ and iAKO mice. n=10-11.

## References

1. S. Yu et al., Adipocyte-specific gene expression and adipogenic steatosis in the mouse liver due to peroxisome proliferator-activated receptor gamma1 (PPARgamma1) overexpression. The Journal of biological chemistry 278, 498–505 (2003).

2. Y. H. Hong et al., Up-regulation of adipogenin, an adipocyte plasma transmembrane protein, during adipogenesis. Molecular and cellular biochemistry 276, 133–141 (2005).

3. J. Y. Kim, K. Tillison, C. M. Smas, Cloning, expression, and differentiation-dependent regulation of SMAF1 in adipogenesis. Biochemical and biophysical research communications 326, 36–44 (2005).

4. G. Ren, P. Eskandari, S. Wang, C. M. Smas, Expression, regulation and functional assessment of the 80 amino acid Small Adipocyte Factor 1 (Smaf1) protein in adipocytes. Archives of biochemistry and biophysics 590, 27–36 (2016).

5. T. O. Kilpeläinen et al., Genome-wide meta-analysis uncovers novel loci influencing circulating leptin levels. Nature communications 7, 10494 (2016).

6. A. Alvarez-Guaita et al., Phenotypic characterization of Adig null mice suggests roles for adipogenin in the regulation of fat mass accrual and leptin secretion. Cell reports 34, 108810 (2021).

7. A. J. Mathiowetz, J. A. Olzmann, Lipid droplets and cellular lipid flux. Nature cell biology 26, 331–345 (2024).

8. M. J. Rao, J. M. Goodman, Seipin: harvesting fat and keeping adipocytes healthy. Trends in cell biology 31, 912–923 (2021).

9. W. Fei et al., Fld1p, a functional homologue of human seipin, regulates the size of lipid droplets in yeast. The Journal of cell biology 180, 473–482 (2008).

10. K. M. Szymanski et al., The lipodystrophy protein seipin is found at endoplasmic reticulum lipid droplet junctions and is important for droplet morphology. Proceedings of the National Academy of Sciences of the United States of America 104, 20890–20895 (2007).

11. Z. Cao, Y. Hao, C. W. Fung, Dietary fatty acids promote lipid droplet diversity through seipin enrichment in an ER subdomain. Nature communications 10, 2902 (2019).

12. Y. Tian et al., Tissue-autonomous function of Drosophila seipin in preventing ectopic lipid droplet formation. PLoS genetics 7, e1001364 (2011).

13. L. Liu et al., Adipose-specific knockout of SEIPIN/BSCL2 results in progressive lipodystrophy. Diabetes 63, 2320–2331 (2014).

14. X. Cui et al., Seipin ablation in mice results in severe generalized lipodystrophy. Human molecular genetics 20, 3022–3030 (2011).

15. J. Magré et al., Identification of the gene altered in Berardinelli-Seip congenital lipodystrophy on chromosome 11q13. Nature genetics 28, 365–370 (2001).

16. H. Wang et al., Seipin is required for converting nascent to mature lipid droplets. eLife 5, (2016).

17. V. T. Salo, I. Belevich, S. Li, Seipin regulates ER-lipid droplet contacts and cargo delivery. The EMBO journal 35, 2699–2716 (2016).

18. V. T. Salo et al., Seipin Facilitates Triglyceride Flow to Lipid Droplet and Counteracts Droplet Ripening via Endoplasmic Reticulum Contact. Developmental cell 50, 478–493.e479 (2019).

19. H. Arlt et al., Seipin forms a flexible cage at lipid droplet formation sites. Nature structural & molecular biology 29, 194–202 (2022).

20. R. Yan et al., Human SEIPIN Binds Anionic Phospholipids. Developmental cell 47, 248–256.e244 (2018).

21. C. Wang, A Sensitive and Quantitative mKeima Assay for Mitophagy via FACS. Current protocols in cell biology 86, e99 (2020).

22. J. D. Martell, T. J. Deerinck, S. S. Lam, M. H. Ellisman, A. Y. Ting, Electron microscopy using the genetically encoded APEX2 tag in cultured mammalian cells. Nature protocols 12, 1792–1816 (2017).

23. H. Lorenz, D. W. Hailey, C. Wunder, J. Lippincott-Schwartz, The fluorescence protease protection (FPP) assay to determine protein localization and membrane topology. Nature protocols 1, 276–279 (2006).

24. J. Chen, A. D. Brunner, Pervasive functional translation of noncanonical human open reading frames. Science (New York, N.Y.) 367, 1140–1146 (2020).

25. D. M. Anderson et al., A micropeptide encoded by a putative long noncoding RNA regulates muscle performance. Cell 160, 595–606 (2015).

26. J. Chung et al., LDAF1 and Seipin Form a Lipid Droplet Assembly Complex. Developmental cell 51, 551–563.e557 (2019).

27. I. G. Castro et al., Promethin Is a Conserved Seipin Partner Protein. Cells 8, (2019).

28. Y. A. Klug, J. C. Deme, R. A. Corey, M. F. Renne, Mechanism of lipid droplet formation by the yeast Sei1/Ldb16 Seipin complex. Nature communications 12, 5892 (2021).

29. D. Binns, S. Lee, C. L. Hilton, Q. X. Jiang, J. M. Goodman, Seipin is a discrete homooligomer. Biochemistry 49, 10747–10755 (2010).

30. X. Sui et al., Cryo-electron microscopy structure of the lipid droplet-formation protein seipin. The Journal of cell biology 217, 4080–4091 (2018).

31. B. R. Cartwright, J. M. Goodman, Seipin: from human disease to molecular mechanism. Journal of lipid research 53, 1042–1055 (2012).

32. C. Windpassinger et al., Heterozygous missense mutations in BSCL2 are associated with distal hereditary motor neuropathy and Silver syndrome. Nature genetics 36, 271–276 (2004).

33. V. A. Payne et al., The human lipodystrophy gene BSCL2/seipin may be essential for normal adipocyte differentiation. Diabetes 57, 2055–2060 (2008).

34. W. Fei et al., Molecular characterization of seipin and its mutants: implications for seipin in triacylglycerol synthesis. Journal of lipid research 52, 2136–2147 (2011).

35. J. Abramson et al., Accurate structure prediction of biomolecular interactions with AlphaFold 3. Nature, (2024).

36. X. Prasanna et al., Seipin traps triacylglycerols to facilitate their nanoscale clustering in the endoplasmic reticulum membrane. PLoS Biol 19, e3000998 (2021).

37. A. Bartelt et al., Brown adipose tissue activity controls triglyceride clearance. Nat Med 17, 200–205 (2011).

38. P. P. Khedoe et al., Brown adipose tissue takes up plasma triglycerides mostly after lipolysis. Journal of lipid research 56, 51–59 (2015).

39. H. Zhou, C. Xu, H. Lee, Y. Yoon, W. Chen, Berardinelli-Seip congenital lipodystrophy 2/SEIPIN determines brown adipose tissue maintenance and thermogenic programing. Mol Metab 36, 100971 (2020).

40. L. Dollet et al., Seipin deficiency alters brown adipose tissue thermogenesis and insulin sensitivity in a non-cell autonomous mode. Sci Rep 6, 35487 (2016).

41. S. V. Kathuria, Y. H. Chan, R. P. Nobrega, A. Özen, C. R. Matthews, Clusters of isoleucine, leucine, and valine side chains define cores of stability in high-energy states of globular proteins: Sequence determinants of structure and stability. Protein Sci 25, 662–675 (2016).

42. T. C. Walther, S. Kim, H. Arlt, G. A. Voth, R. V. Farese, Jr., Structure and function of lipid droplet assembly complexes. Curr Opin Struct Biol 80, 102606 (2023).

43. V. Zoni et al., Seipin accumulates and traps diacylglycerols and triglycerides in its ring-like structure. Proceedings of the National Academy of Sciences of the United States of America 118, (2021).

44. S. Kim et al., Seipin transmembrane segments critically function in triglyceride nucleation and lipid droplet budding from the membrane. Elife 11, (2022).

45. M. Jiang et al., Lack of testicular seipin causes teratozoospermia syndrome in men. Proceedings of the National Academy of Sciences of the United States of America 111, 7054–7059 (2014).

46. D. Ito, N. Suzuki, Seipinopathy: a novel endoplasmic reticulum stress-associated disease. Brain 132, 8–15 (2009).

47. W. Fei et al., A role for phosphatidic acid in the formation of "supersized" lipid droplets. PLoS genetics 7, e1002201 (2011).

48. J. Bi et al., Seipin promotes adipose tissue fat storage through the ER Ca²⁺-ATPase SERCA. Cell Metab 19, 861–871 (2014).

49. Y. Combot et al., Seipin localizes at endoplasmic-reticulum-mitochondria contact sites to control mitochondrial calcium import and metabolism in adipocytes. Cell reports 38, 110213 (2022).

